# How to scale up from animal movement decisions to spatio-temporal patterns: an approach via step selection

**DOI:** 10.1101/2022.07.19.500568

**Authors:** Jonathan R. Potts, Luca Börger

## Abstract

1. Uncovering the mechanisms behind animal space use patterns is of vital importance for predictive ecology, thus conservation and management of ecosystems. Movement is a core driver of those patterns so understanding how movement mechanisms give rise to space use patterns has become an increasingly active area of research.
2. This study focuses on a particular strand of research in this area, based around step selection analysis (SSA). SSA is a popular way of inferring drivers of movement decisions, but, perhaps less well-appreciated, it also parametrises a model of animal movement. Of key interest is that this model can be propogated forwards in time to predict the space use patterns over broader spatial and temporal scales than those that pertain to the proximate movement decisions of animals.
3. Here, we provide a guide for understanding and using the various existing techniques for scaling-up step selection models to predict broad scale space use patterns. We give practical guidance on when to use which technique, as well as specific examples together with code in `R` and `Python`.
4. By pulling together various disparate techniques into one place, and providing code and instructions in simple examples, we hope to highlight the importance of these techniques and make them accessible to a wider range of ecologists, ultimately helping expand the usefulness of step selection analysis.

## 1 Introduction

Understanding and predicting the spatio-temporal distribution of individuals, populations and communities is a fundamental aim of ecological research. Key drivers of these spatial dynamics are the movement decisions of individuals in response to individual and local environmental conditions and resources (Nathan *et al.*, 2008; Fryxell *et al.*, 2008), the distribution of conspecific and heterospecific individuals (Osborne *et al.*, 2022) and environmental change (Tuomainen & Candolin, 2011) and disturbance (Courbin *et al.*, 2022). Animal movements fundamentally affect other ecological processes, including population (Hamilton & May, 1977) and community dynamics (Costa-Pereira *et al.*, 2022), transport processes (Abbas *et al.*, 2012), disease spread (Merkle *et al.*, 2018), and ecosystem processes (Doughty *et al.*, 2016). Under current global change it is becoming increasingly important to robustly predict changes in individual movement decisions and how these scale up to emergent spatial patterns of animal distributions.

Focusing on environmental conditions and resources, habitat selection methods aim to identify and quantify the link between animal movements and distributions and the environment (Fieberg *et al.*, 2021) and are fundamentally based on a comparison between the distribution of environmental resources and the proportional use of these by the individuals (Manly *et al.*, 2002). A key limitation of such approaches lies in the definition of what is available to each individual (Buskirk & Millspaugh, 2006), as animals are fundamentally limited by their movement capacities and cannot reach every point in the landscape at every single movement step, an issue which has led long-standing discussions and methodological debates in the literature (McClean *et al.*, 1998; Northrup *et al.*, 2013). This crucial limitation can be solved by integrated step-selection analysis (iSSA), which allows simultaneous modelling of movement and habitat selection decisions by animals (Avgar *et al.*, 2016), building upon the earlier technique of step selection analysis (SSA) (Fortin *et al.*, 2005; Rhodes *et al.*, 2005; Forester *et al.*, 2009; Thurfjell *et al.*, 2014). Not only has this fundamental methodological advancement lead to an explosion of the use of SSA and iSSA in recent years (Viana *et al.*, 2018; Huggler *et al.*, 2022; Northrup *et al.*, 2022), and methodological extensions (Munden *et al.*, 2021; Klappstein *et al.*, 2021), but researchers have increasingly shown how the movement kernels parameterised during SSA can be ‘scaled-up’ to predict broader-scale space use patterns (Potts *et al.*, 2014b; Avgar *et al.*, 2016; Signer *et al.*, 2017; Potts & Schlägel, 2020; Fieberg *et al.*, 2021). Even though this markedly increases the level of understanding and the quality of predictions which can be obtained from animal movement analyses, such upscaling of step selection analysis is seldom done by the many studies using SSA, perhaps due to a lack of knowledge or perceived methodological difficulties.

The focus of this methods guide is to elucidate the various methods for scaling up from step selection to utilisation distributions and other broader space use patterns. We aim to guide the reader through the various techniques used for scaling up, when they can be applied and when not, and summarising the pros and cons when more than one technique is potentially applicable. We also give example code, in both `R` and `Python`, of some simple case studies, to give the reader a practical way of getting started with these techniques.

Throughout, we will distinguish between two different types of movement models, which each require slightly different methods of analysis. The first type models the movement decisions animals make due to spatial variables that remain essentially unaffected by the animals’ presence (e.g. terrain, weather). In these situations, one can use a correlative model, whereby the animal movement decision is the response variable and the unaffected spatial variable is the explanatory variable, and propogate that model forwards through time, as exemplified by studies such as (Potts *et al.*, 2014a; Signer *et al.*, 2017).

In the second type of movement model, there are variables that both affect an animal’s movement and are affected by the same animal’s presence. Examples include between-animal interactions, whereby the presence of individual A (either in the present or recent past) affects the movement of individual B, but in turn the movements of individual A are affected by the presence individual B (Couzin *et al.*, 2002; Giuggioli *et al.*, 2013). Another example is where animal movement is affected by the presence or absence of some resource that they then consume and deplete. Thus these animals also affect the resource landscape by their presence, creating a feedback loop between animal location and landscape variable (Riotte-Lambert & Matthiopoulos, 2020), which may also be mediated by memory (Lewis *et al.*, 2021). In all these situations, whilst it is possible to use correlative models to make inferrence about the effect of variable U on variable V (e.g. the presence of individual A on the movement of individual B), these correlative models cannot be reliably propogated forwards in time to predict broad space use patterns, since the reality is that both variables affect each other. There is no *a priori* sense in which one is the explanatory variable and the other the response variable. Instead, scaling up to broader spatio-temporal scales requires a dynamic modelling approach, for example via individual based models (IBMs) of interacting individuals (Avgar *et al.*, 2013) or systems of partial differential equations (PDEs) (Moorcroft & Lewis, 2006; Potts & Lewis, 2019).

Analysing these dynamic models is much more complicated than the case of purely correlative models, and we cannot give a complete hands-on guide here. Our approach will therefore be to provide a gateway into these techniques by explaining how to construct such IBMs and PDEs from the output of step selection, and pointing to the possible methods of analysis that might be used. In the case of IBMs, we do provide some code to help the reader get started. However, in the case of PDE models, there is a world of analytic techniques, which are standard tools for many applied mathematicians, but may not be familiar to those without an applied mathematics background. Our guide on PDE techniques will not be to cover that vast background itself, but rather to *guide collaborations* between ecologists and applied mathematicians by showing how to interface questions about spatial arrangements of ecological systems with mathematical tools for deriving emergent spatial phenomena. We contend that such inter-disciplinary collaborations are perhaps the only way forwards for answering many important questions in spatial ecology.

The ‘scaling up’ procedure we will describe is outlined as follows. The first step is to parametrise a step selection function from empirical data. This step is typically called step selection analysis. A recent ‘How to’ paper already exists that covers the practicalities and interpretation of step selection analysis (Fieberg *et al.*, 2021), so we will be relatively brief here, referring the interested reader to that paper, also noting the recent review by Northrup et al. (2022). The second step involves using the parametrised step selection function to infer broad-scale space-use patterns. We will describe various existing techniques for this, which use different mathematical formalisms. Some of these techniques give exact results and some are approximate, some stochastic and some deterministic. Moreover, not all of these techniques can be used in every situation. Therefore we will give a guide as to which technique can (or should) be used in which situations.

Finally, we will explain the sort of information one can gain from this approach. This includes (a) predicting whether or not one might expect steady state space-use distributions to emerge (the alternative being those that are in perpetual flux), (b) predicting the shape of such emergent steady-state distributions (e.g. stable home ranges) where they exist, (c) using the emergent spatial patterns as a goodness-of-fit test, which can help detect missing features in step selection models. The methods presented throughout necessarily involve quite a bit of mathematical formalism. To aid the reader in navigating this, we include various supplementary appendices that give specific examples of our mathematical formalisms, as well as code for implementing the methods in these simple examples, in both `R` and `Python`.

## 2 Step selection models of animal movement

The principle aim of step selection analysis (SSA) is to understand the drivers of animal movement. Such movements can be affected by an incredibly wide range of possible phenomena, including food distribution, predator avoidance, social interactions, corridors to ease passage, physical barriers, topography, and many more, together called movement covariates. All of these can potentially be revealed through a step selection approach, as long as one has the right data. SSA is a type of resource selection analysis, i.e. it is a way of estimating the probability that an animal will use a given spatial area, as a function of that area’s ‘resource value’ (e.g. access to food, mates, resting area, thermal refuge, etc.). SSA specifically focuses on the movement between two locations. It compares the observed movement step to all other locations the animal could have reached during that same time step, given the movement capacity of the animal, aiming to determine why the animal took the particular observed step instead of the many available alternatives.

With the increasingly detailed data on animal movement and their covariates that has become available over the past years, SSA has become a very popular approach for inferring drivers of movement. There have also been various variants of SSA introduced, such as integrated SSA (iSSA; Avgar *et al.* (2016)), and time-varying iSSA (tiSSA; Munden *et al.* (2021)). However, for simplicity we will refer to all of these as SSA unless there is a good reason to be specific. To understand how to use SSA for *inference,* there is a recent ‘How to’ paper, which also deals with habitat selection more broadly (Fieberg *et al.*, 2021). Those unfamiliar with SSA may want to read this paper before continuing. Other useful papers related to inferrence in SSA are Thurfjell *et al.* (2014); Northrup *et al.* (2022).

Here, however, our focus is different. Instead of focusing on inference, our aim is to give a guide for how to use the output of SSA for predicting space use patterns, given the information provided by SSA on both an animal’s preferences for resources (i.e. movement covariates) and its movement behaviour. In more formal terms, we start by observing that a by-product of SSA (or iSSA) is the parametrisation of a *movement kernel,* describing the probability density, *p_τ_*(**z**|**x**, *α*_**x**_, *t*) of moving from one location, **x**, to another location, **z**, between times *t* and *t* + *τ*, given all the information we have on the movement covariates, and given that the animal travelled to **x** on a bearing of *α*_**x**_. We want to examine how to derive a *utilisation distribution* from this movement kernel, which is the probability distribution of finding the animal at any given location, and closely related to the concept of ‘home range’ (Börger *et al.*, 2008). This requires setting up quite a bit of mathematical notation, but a picture of what is going on underneath these equations is given in Fig. 1, and we give some foundational examples in Supplementary Appendix A to help the reader get to grips with the mathematical formalisms. We will also focus on location data that are recorded at fixed intervals at a relatively low frequency (e.g. one every few minutes or hours), but see Supplementary Appendix B for modifications away from this case (following Potts *et al.* (2018); Munden *et al.* (2021)).

**Fig. 1.**
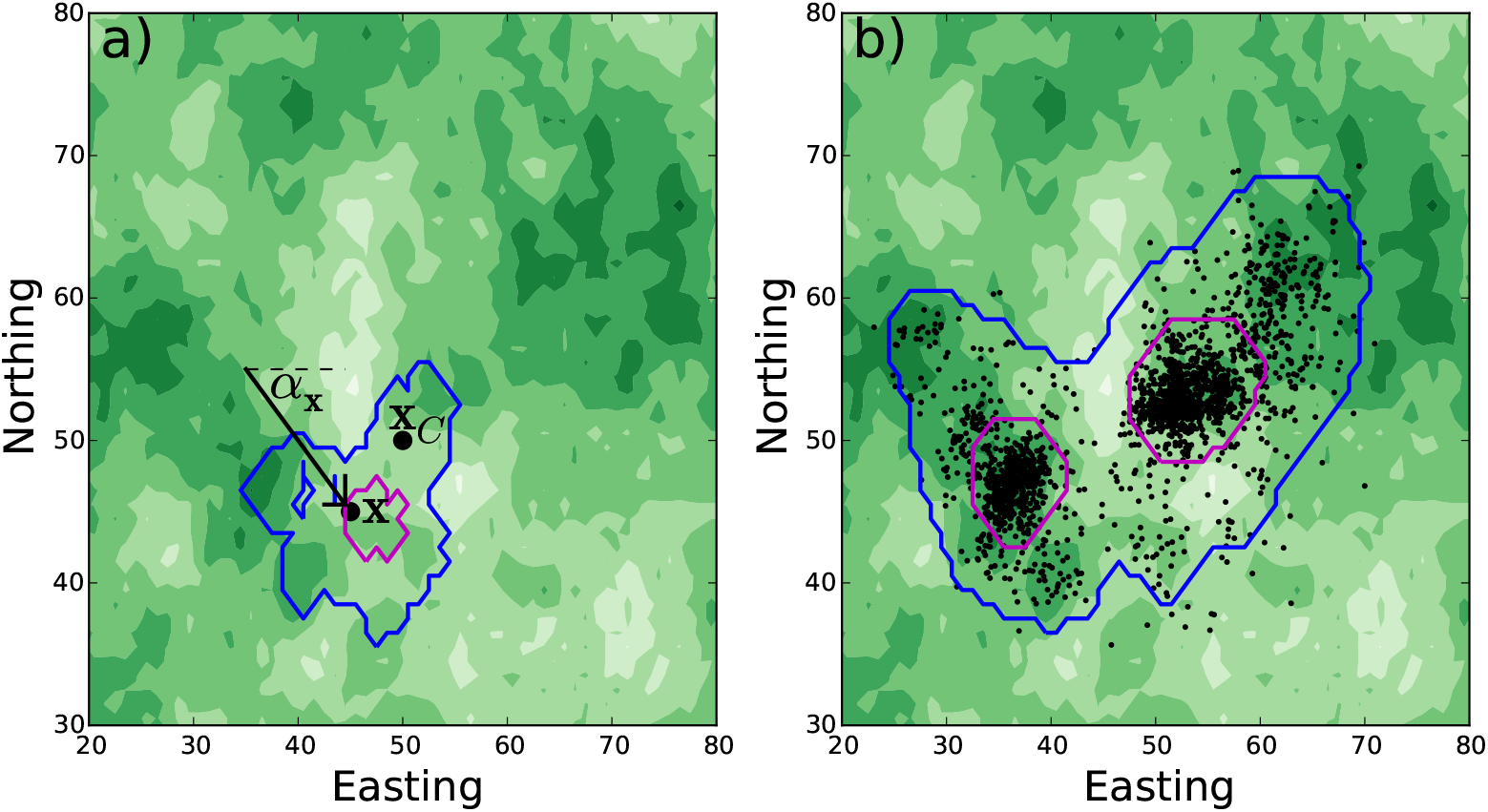
From a movement kernel to a utilisation distribution. Panel (a) shows the movement kernel for a simulated animal that has a biases towards (i) better resources, denoted by darker green, (ii) a central place at **x**_*C*_, and (iii) to continuing in the same direction (i.e. correlated movement). It has arrived at location **x** at time *t* on a bearing of *α*_**x**_. The animal has a 95% (resp. 50%) probability of being located in the blue (resp. magenta) contour at time *t* + *τ* (i.e. after one time step). Panel (b) shows the utilisation distribution after 2000 time steps, where the blue (resp. magenta) curves show 95% (resp. 50%) kernel density estimator, mimicking how home ranges are often calculated from field data. This paper aims to explain various techniques for mathematically deriving, hence predicting, the utilisation distribution (Panel b) from knowledge of the movement kernel (Panel a).

### 2.1 The movement kernel from step selection

Right away, let us write down the general form for our movement kernel, which combines two sets of equations, one to estimate the intrinsic movement behaviour of the animals and one for estimating the selection for resources, and is written as

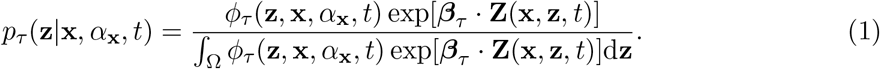

We now explain all the terms from Equation (1) in detail. First, **Z**(**x**, **z**, *t*) is a vector of movement covariates. We write **Z** = (*Z*_1_,…,*Z_n_*) where, for each *i* = 1,…,*n*, the function *Z_i_* = *Z_i_*(**x**, **z**, *t*) may depend upon any combination of the start location **x**, the end location **z**, or the current time *t*. However, not every *Z_i_* need depend upon all of these aspects. For example, if the *i*-th covariate is something constant in time over the duration of the study, e.g. height above sea level, then *Z_i_* would not depend upon *t*. Also, *Z_i_* may depend upon the start location (at time *t*), the end location (at time *t* + *τ*) or both. If *Z_i_* depends on both **x** and **z**, this implicitly means it could depend upon any or all of the points between **x** and **z**.

Second, *β_τ_* is a vector denoting the strength of the effect of each covariate, *Z_i_*. We use the subscript *τ* in Equation (1) to emphasise that *β_τ_* may depend upon *τ* (Fieberg *et al.*, 2021). However, for notational convenience, we will usually drop this subscript and write *β_τ_* = *β* = (*β*_1_,…, *β_n_*). The function exp[*β_τ_*·***Z***(**x**, **z**, *t*)] is sometimes called a step selection function (Fortin *et al.*, 2005), but confusingly this term has also been used synonymously with the movement kernel (Forester *et al.*, 2009; Potts *et al.*, 2014a). Note that we can expand the scalar product of *β_τ_* and **Z**(**x**, **z**, *t*) in Equation (1) to give *β_τ_* · ***Z***(**x**, **z**, *t*) = *β*_1_*Z*_1_(**x**, **z**, *t*) + ⋯ + *β_n_Z_n_*(**x**, **z**, *t*).

Third, *ϕ_τ_*(**z, x**, *α*_x_, *t*) is a selection-free movement kernel, and has historically taken various functional forms, discussed in Fieberg et al. (2021, Section 3.1). The function *ϕ_τ_* is sometimes referred to as a resource-free or resource-independent movement kernel. However, here we will use the term ‘selection-free’, as in principle *ϕ_τ_* could itself depend upon resources and other environmental features (Avgar *et al.*, 2016). This kernel can be thought of as describing the intrinsic ability of an animal to move in a given environment, disregarding any decisions the animal makes about where to move. In the simplest examples, *ϕ_τ_* is just a decaying function of the speed |**z** – **x**|/*τ*, e.g. *ϕ_τ_*(**z**,**x**, *α*_x_, *t*) = exp(-|**z** – **x**|/*τ*), but it can have a more complicated dependence on the landscape and/or time (Avgar *et al.*, 2016). Note that the possible dependence on *α*_x_ allows for *ϕ_τ_* to incorporate a distribution of turning angles, allowing for correlated movement.

Finally, Ω is the study area and the denominator in Equation (1) ensures that the movement kernel integrates to 1 with respect to **z**, making *p_τ_*(**z**|**x**, *α*_x_, *t*) a genuine probability density function. It is sometimes convenient to write

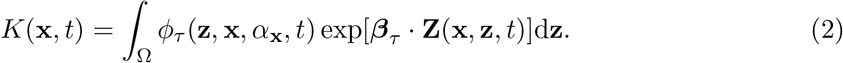

Equation (1) appears as Equation (13) in Fieberg *et al.* (2021) in a slightly different form. We explain in Supplementary Appendix A how to relate the two formulae precisely, to enable smooth linkage between both papers, alongside some examples of movement kernels with code.

### 2.2 Incorporating animal interactions

Whilst Equation (1) models the effect of covariates on an animal’s movement, some covariates are also affected by the animal’s movement. This means the two features of interest, animal locations and covariate values, feedback on one another. A key example is when an animal’s movement is affected by the space use of a second animal, whose movement is in turn affected by the space use of the first animal. Such two-way interactions may be social, competitive, mutualistic, or predator-prey. In any such case, SSA will lead to a different movement kernel for each animal, which interact with each other. This situation leads to a *system* of *coupled* movement kernels, sometimes called ‘coupled step selection functions’ (Potts *et al.*, 2014b; Lewis *et al.*, 2021). If we have *N* interacting animals then the general form of such a system is

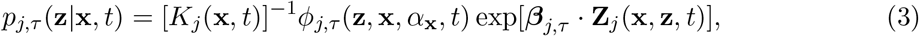

where *j* = 1,…, *N* denotes the number of the animal.

An example system of coupled movement kernels is where each animal j has at least one covariate *Z_i,j_* (for some *i* between 1 and *n*) that denotes the recent space use of a different animal, say animal *j*′ where *j*′ = *j* (perhaps mediated through terrain marking or memory (Moorcroft & Lewis, 2006; Riotte-Lambert *et al.*, 2015)). This means that the movement of animal *j* is affected by the locations of animal *j*′. But if, in turn, animal *j*′ has a covariate, say *Z*_*i′, j′*_, that denotes the space use of animal *j*, then the two animal’s movements depend on one another, generating a feedback loop between them. This is what we mean by saying the equations are ‘coupled’.

In principle, Equation (3) can be parametrised using SSA in exactly the same way as Equation (1), for each individual *j* = 1,…, *N*. However, more needs to be said to put this into practice, as it is not a trivial task to determine the ‘recent space use’ of each animal. Indeed, even conceptually the idea of ‘recent space use’ begs questions. How recent? What constitutes ‘space use’ ? The precise one-dimensional path the animal has travelled or some broader area demarcated by that path? How do we infer this area from data?

None of these questions have a single catch-all answer. However, an important step forward was made by Schlägel *et al.* (2019). Their method starts by using the notion of an *occurrence distribution* (OD) to describe the ‘recent space use’ of an animal (Fleming *et al.*, 2016). The OD is constructed by taking consecutive measured locations of an animal, over a user-defined timeframe, and building a theoretically optimal estimation of the distribution of actual locations. The `R` package `ctmm` enables users to construct this OD with just a few lines of code, as explained in Calabrese *et al.* (2016). However, the output can be quite finely resolved, and in practice animals may respond to a more smoothed version of the OD. Therefore it is worth users also experimenting with using either a spatial averaging of the OD, or something more instrinsically smoothed-out, like KDE (Worton, 1989) or AKDE (Fleming *et al.*, 2015).

Whichever method is chosen, to use the OD for SSA simply involves considering the OD in exactly the same way as any other environmental layer that varies through time. To put this in mathematical notation, let us denote the value of the OD of animal *j* at time *t* and location *ξ* by *O_j,t_*(*ξ*) (where *ξ* stands for either **x** or **z**). If we hypothesise that this OD covaries with the decision of an animal *j*′ to move *to* a location then we set *Z_i,j′_*(**x**, **z**, *t*) = *O_j,t_*(**z**) for some *i* (Schlägel *et al.*, 2019). Alternatively, if we hypothesise that *O_j,t_*(*ξ*) covaries with the decision of *j*′ to move *from* a location then we set *Z_i,j′_*(**x**, **z**, *t*) = *O_j,t_*(**x**) for some *i*. Following this procedure for each pair of animals *j, j′* = 1,…, *N* leads to a collection of covariates in a system of coupled movement kernels (Equation 3), which can be parametrised using SSA (Avgar *et al.*, 2016; Fieberg *et al.*, 2021).

## 3 Emergent patterns from scaling up

We now turn to the main topic of this paper, which is how to scale up from the movement kernel of Equation (1) to a description of broad scale space use patterns. For this, we use the concept of a utilisation distribution (UD), which measures the probability of finding an animal at a location **x** at time t. We denote the UD by *u*(**x**, *t*). We will sometimes be interested in the UD as it changes over time, but we will also examine situations where it is possible to derive a steady state UD, *u*_*_(**x**), which denotes the limit as *t* → ∞ of *u*(**x**, *t*). Mathematically, it is not always the case that *u*_*_(**x**) exists, as *u*(**x**, *t*) may exhibit oscillatory behaviour at long times, or more complicated spatio-temporal fluctuations (Potts & Lewis, 2019). However, where it does exists, and where this limit is approached in an ecologically-relevant time frame, the UD corresponds to what is usually called a home range.

In this section, we will examine how to derive both exact and approximate expressions for *u*(**x**, *t*) from Equation (1). We will explain the situations in which one can use each of these expressions, and the various benefits and drawbacks of each. In general, whilst exact expressions are theoretically ideal, they may be either difficult to compute in practice, not amenable to mathematical analysis, or only apply to certain subcases of Equation (1). Approximate expressions are therefore also valuable. We will explain how to calculate each of the expressions in practice and also provide a gateway into mathematical analysis. Finally, we will explain how to determine whether *u*_*_(**x**) exists or not, and how to compute it where possible.

### 3.1 Mathematical formalisms for scaling-up: IDEs, PDEs, IBMs

Here we describe how to use mathematical methods to derive the space use distribution of an animal, such as a home range, starting from the SSA-estimated movement kernel (combining the movement capacity of the animal and the selection for resources in the external environment). We then show how it is then possible to predict the space use distribution of an animal in a landscape with a certain distribution of resources, based on knowledge of its step-to-step movement decisions.

Perhaps the most general form linking a movement kernel to a UD is the so-called *master equation* (van Kampen, 1981; Merkle *et al.*, 2017), which is an example of an integro-difference equation (IDE). We will begin by assuming that *p_τ_* is independent of *α*_x_ (i.e. no correlation in movement), so that *p_τ_*(**z**|**x**, *α*_x_) = *p_τ_*(**z**|**x**, *t*). The master equation in this case is

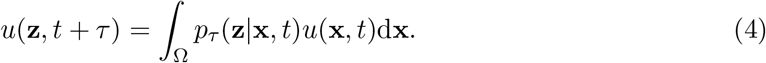

Intuitively, this equation says

1. start with a utilisation distribution at time *t*, given by *u*(**x**, *t*),
2. then multiply this by the probability density of moving from **z** to **x** in a timestep of length *τ*, given by *p_τ_*(**z**|**x**, *t*),
3. do this for all **x** and integrate,
4. and then the result is the probability distribution at time *t* + *τ*.

It is also possible to incorporate the aspect of correlated movement (i.e. dependence on *α*_x_) with some extra notational baggage. We explain this in Supplementary Appendix C, but focus here on uncorrelated movement for ease of explanation.

One approach to calculating *u*(**x**, *t*) is to solve Equation (4) numerically. In practice, this involves discretising the study region, Ω, and turning the integral into a sum. Let us denote by *S* a set of points obtained from discretising Ω. This discretisation can be chosen by the user, but in practice it makes sense to use a grid that is related to the underlying environmental covariates, as these themselves often arrive as a discrete-space raster. Then, for any pair of grid cells, *s* and *s*′, write the probability of moving from *s*′ to *s* as *P_τ_*(*s*|*s*′, *t*).

Next let *U*(*s, t*) denote the probability of finding an animal at grid-point *s* at time *t*. Then, given this discretisation, Equation (4) becomes

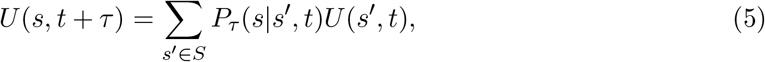

An example of calculating Equation (5) over time is given in Supplementary Appendix D, together with code.

Equation (4) gives an exact solution for the time-evolution of the UD, given a movement kernel. Therefore, in principle, it gives a complete description of how to scale up from a movement kernel to a UD. Furthermore, Equation (5) gives a way of calculating (4), with only some minor approximations due to the discretisation. So it might feel like the job is done. However, there are two key downsides to these equations. The first is that they are not particularly amenable to exact mathematical analysis, so there is not much exact theory that one can draw on (but see Barnett & Moorcroft (2008), which we discuss in Section 3.2). The second is that a numerical solution can be very time-consuming. Calculating Equation (5) requires calculation of *P_τ_*(*s*|*s*′, *t*) for every *s, s*′ ∈ *S*, and *P_τ_*(*s*|*s*′, *t*) itself requires computational of a numerical integral, the demoninator in Equation (1). We now deal with each drawback in turn and how to mitigate against them.

First, to make use of mathematical theory, it is beneficial to derive a partial differential equation (PDE) from Equation (4), as PDEs are in general far more amenable to mathematical analysis than IDEs. However, it does require assumptions to be made. Potts & Schlägel (2020) examined the PDE limit in a situation where we assume the master equation is of the form in Equation (4). Additionally, they made two assumptions about the movement kernel in Equation (1). First, *ϕ_τ_* must be function of only |**z** – **x**|, so that *ϕ_τ_*(**x**, **z**, *α*_x_, *t*) = *ψ_τ_*(|**x** – **z**|). Second, **Z** must be function of only **z** (the end of the step) and t. Third, *τ* must be sufficiently small so that one cannot reasonably expect an animal to have made several decisions about where to move during this time (e.g. if the landscape over which the animal roams during a single step of time *τ* fluctuates greatly, causing multiple turns). Given these, Potts & Schlägel (2020) showed that the UD is approximately governed by the following PDE

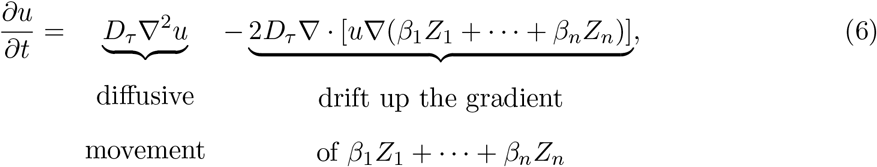

where *D_τ_* is the diffusion constant, calculated as

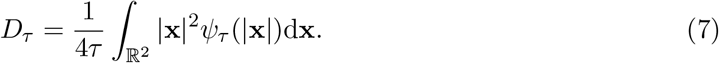

From the perspective of calculating *u*(**x**, *t*) numerically, there is in our experience no advantage to using this PDE over the master equation. However, the analytic tools for studying PDEs are vast and can give crucial qualitative information about space use, which we will discuss more in Section 4.

As an alternative to numerical solutions of either the PDE in Equation (6) or the IDE in Equation (4), a conceptually-simple approach is to use a stochastic individual based model (IBM) (Avgar *et al.*, 2016; Signer *et al.*, 2017; Potts *et al.*, 2022a). This simply involves simulating stochastic realisations of the movement kernel in Equation (1) and can be done either directly (Supplementary Appendix A) or via the `amt` package in `R`. The `amt` approach is shown, with examples, in Signer *et al.* (2019). Here, we only want to add one word of caution, which is that the UD described in Signer *et al.* (2019), and calculated in `amt`, is in fact the time-integral of *u*(**x**, *t*), i.e.

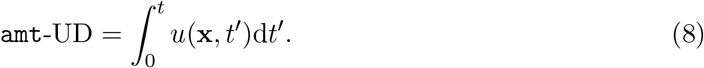

If the UD reaches a steady state, *u*_*_(**x**), then the two definitions of UD coincide, but this is not true for transient UDs. Therefore anyone comparing transient UDs from `amt` with those calculated from the IDE and PDE methods described here needs to take this discrepancy into account. In the context of home range calculations, the `amt` method is closer to what is done in practice when measuring home ranges from data, as typically people will measure home ranges using highly autocorrelated movement data. However, conceptually, a home range is usually thought of as reflecting the probability density function of an animal’s locations, which is more accurately described by *u*(**x**, *t*).

Whilst IBMs are conceptually simple, it can be time consuming to use them for calculating *u*(**x**, *t*). For this, it is necessary to simulate the movement kernel up to time *t* sufficiently many times to obtain a good measure of the probability distribution, which is the distribution of locations at time *t* across all simulation runs. The number of simulations scales linearly with the number of lattice sites in the study area, and usually one would need dozens or hundreds of simulations per lattice site to obtain a reasonable estimate of the probability distribution. To circumvent this, `amt` estimates Equation (8) instead of *u*(**x**, *t*), and in so doing makes use of every point the simulated animal visits, not just the point at time *t* for each simulation. However, information is lost by estimating Equation (8) rather than *u*(**x**, *t*). Particularly, if *u*(**x**, *t*) fluctuates over time then these fluctuations are averaged-out in the calculation of Equation (8), so are not properly captured by the `amt` method.

If you do want to calculate *u*(**x**, *t*) using IBMs but without requiring prohibitively intensive simulations, it is instead possible to combine simulations with a smoothing method, like kernel density estimation (KDE) (Potts *et al.*, 2014b). To do this, simulate the movement kernel perhaps a few hundred times, each time starting at the same location and running the simulation to time *t* (this much can be done in `amt`). Then take the end-point of each simulation to give a dataset of genuinely independent samples of *u*(**x**, *t*). Finally, construct the KDE of this dataset, which is an estimation of *u*(**x**, *t*).

### 3.2 The steady state UD in the absence of animal interactions

One of the most oft-studied emergent spatial patterns from animal movement data is the home range, i.e. the space use distribution often observed in nature, where animals restrict their movements over time to a certain area in space, and do not roam over the entire available landscape (Börger *et al.*, 2008). Mathematically, this can be thought of as the *steady state* of a UD, defined to be a configuration that does not change over time. As such, if *u*(**x**, *t*) satisfies Equation (4), then the steady state of *u*(**x**, *t*) is a function, *u*_*_(**x**), satisfying the following equation

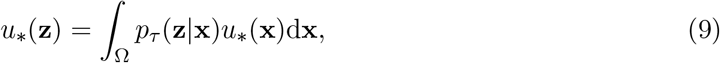

if such a function exists. Here we have to assume that *p_τ_*(**z**|**x**) = *p_τ_*(**z**|**x**, *t*), i.e. the movement kernel does not change over time, otherwise no steady state can exist.

In this section, we will assume that *p_τ_*(**z**|**x**) is independent of *u*(**x**, *t*), i.e. the probability of moving to a specific location does not depend upon the past or present utilisation distribution. This means that Equation (9) is *linear* in *u*_*_(**x**), which is a requirement for the techniques presented in this section. Notice that this linearity requirement precludes the case where we have multiple coupled movement kernels, like in Equation (3) (whereby the movement of one animal depends on the UD of another, whose movement depends on the UD of the first animal). We will return to this case in Section 3.3.

Let us now write the discrete space version of Equation (9), following the notation of Equation (5), as

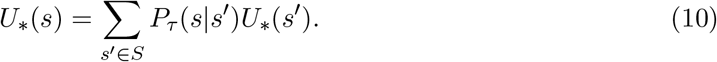

As described in Section 3.1, Equation (10) is what we tend to calculate in practice. This equation can, in theory, be calculated exactly using matrix inversion (Supplementary Appendix E) but to our knowledge this has never been done in the context of step selection, perhaps due to computational intensity.

Another exact method, which is also relatively computationally efficient, is that of Barnett & Moorcroft (2008). This, however, relies on two assumptions. The first is that *ϕ_τ_*(**z**, **x**, *α*_x_, *t*) can be written as a function of |**x** – **z**|, i.e. *ϕ_τ_*(**z**,**x**, *α*_x_, *t*) = *ψ_τ_*(|**x** – **z**|). The second is that the functions *Z_i_*(**x**, **z**, *t*) only depend upon the end-point of the step, i.e. *Z_i_*(**x**, **z**, *t*) = *Z_i_*(**z**) for some function *Z_i_*(**z**). With these assumptions in place, Barnett & Moorcroft (2008) show that the following exact expression for *u\** (**x**) holds

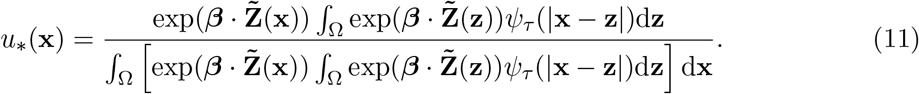

We show how to compute examples of this, with code, in Supplementary Appendix F.

It is interesting to look at two limiting cases. First, if *ψ_τ_* is a uniform distribution, meaning that animals can move over the whole of Ω in a single timestep, then (Supplementary Appendix F)

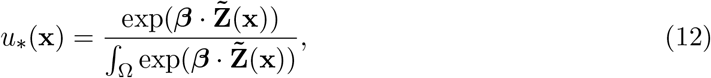

which is just the usual expression for a resource selection function (RSF) (Manly *et al.*, 2002). Although this assumption on *ψ_τ_*(*l*) is quite restrictive, there are real examples. For example, an urban fox can often traverse its whole territory in just a few minutes (Potts *et al.*, 2013), so if Ω were the territory of an urban fox then it makes sense to use a uniform distribution for *ψ_τ_*.

The other extreme is where *ψ_τ_* is arbitrarily narrow, so the animal is making distinct movement choices over much smaller spatial scales than Ω, as is often the case with animals with very large home ranges. In this case we have (Supplementary Appendix F)

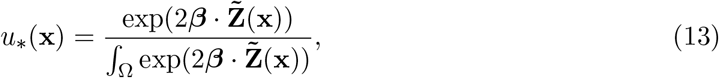

which is identical to Equation (12) but with *β* switched for 2*β*. In other words, the effect of selection on space use doubles as one moves from selection on a very broad spatial scale to a very narrow spatial scale (Moorcroft & Barnett, 2008). Notice that this factor of 2 also appears in the PDE from Equation (6) (before *D_τ_*); indeed, the steady state of Equation (6) is precisely Equation (13) (Potts & Schlägel, 2020).

Figure 2 gives an example of the steady state UD estimations from Equations (11), (12), and (13) for a movement kernel of the following type

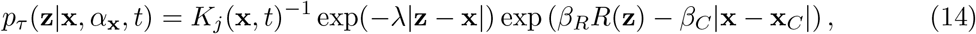

where *R*(**z**) is a resource layer and **x**_*C*_ is a localising point (e.g. a den or nest site). This models movement in a heterogeneous environment where there is additionally some localising tendency towards a single point, such as a den or nest site. Observe from Figure 2 that Equation (12) overestimates the UD size but Equation (13) is an underestimation. We give instructions and example code for reproducing Figure 2 and calculating Equations (11)-(13) in Supplementary Appendix F.

**Fig. 2.**
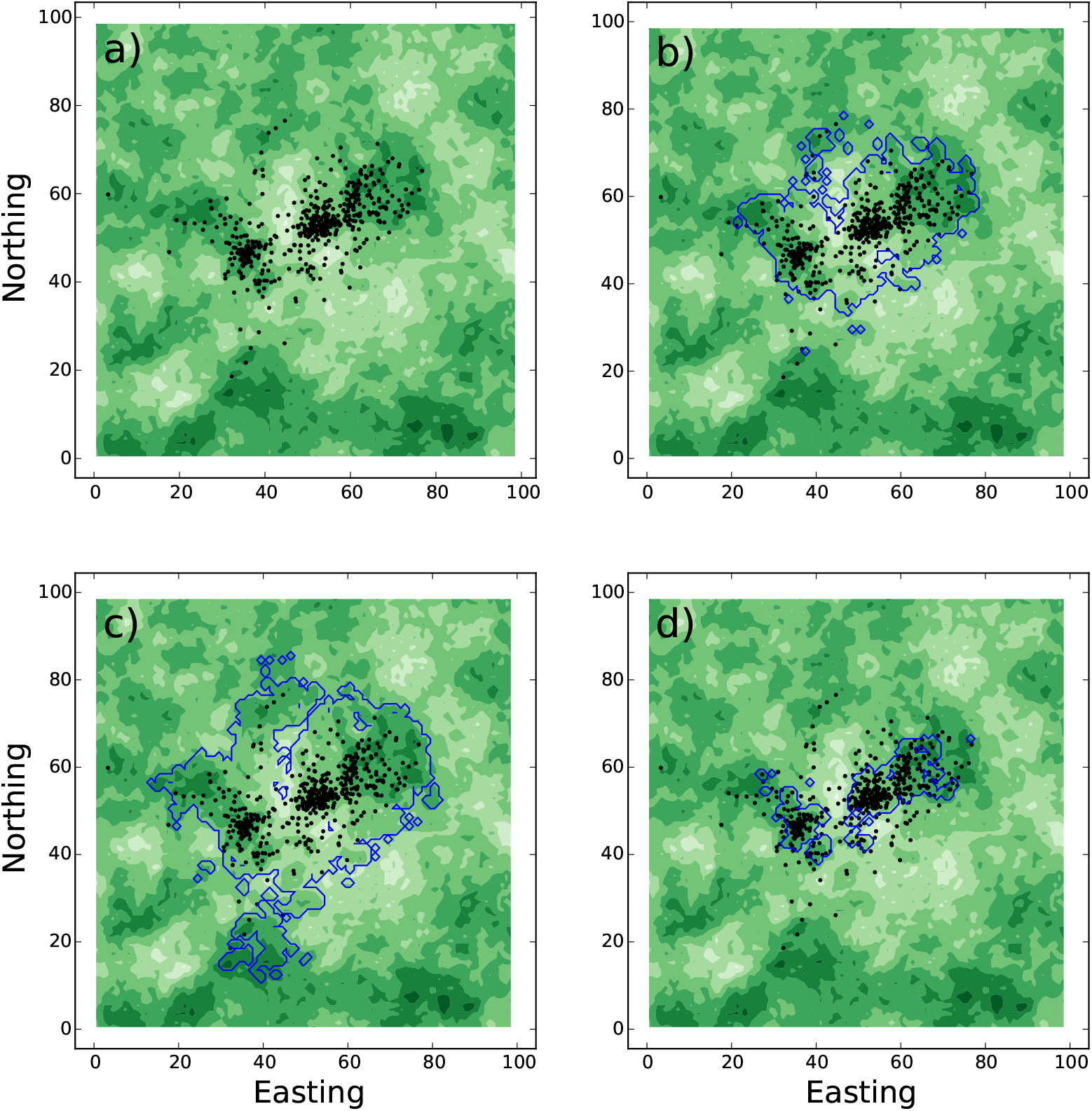
Steady state UDs. Panel (a) shows locations of a simulated animal with movement process corresponding to Equation (14) with the resource layer shown in the background (darker green means higher quality resources). Panels (b-d) overlay this with predicted UDs from Equations (11-13), respectively, shown as a contour line surrounding the 95% kernel of the UD estimation. Panel (b) is an exact steady-state solution to Equation (8), so captures the space use well, whereas Panels (c) and (d) respectively over-and under-estimate space use. Here, λ = 0.2, β_*C*_ = 0.2, and *β_R_* = 1.5, **x**_*C*_ = (50, 50).

### 3.3 Utilisation distributions for interacting animals

In Section 3.2, we examined how to find the steady state distribution in situations where the covariates *Z_i_* are not affected by the locations of the animal or animals. In other words the causality goes one way: covariates affect animal locations, but are not affected by those locations. This works fine in many classical examples of step selection, where the focus is on things like presence of food, ease of motion on the terrain, proximity to locations of interest and so forth. However, in reality there are many situations where movement covariates are affected by present or past animal locations. Examples include memory effects (Merkle *et al.*, 2017), resource depletion (Riotte-Lambert *et al.*, 2015), social interactions (Moorcroft & Lewis, 2006), competition (Vanak *et al.*, 2013), prey-taxis (Kareiva & Odell, 1987), and predator avoidance (Bastille-Rousseau *et al.*, 2015), which are all well-established ecological phenomena.

In such situations, the techniques of Section 3.2 do not directly apply. Indeed, there is no guarantee that a steady state emerges at all, and one may instead find UDs that fluctuate in perpetuity, never settling (Potts *et al.*, 2022b). To understand these patterns, there are two broad approaches: numerical and analytic. For the numerical approach, one could use the IDE formalism from Equation (1) or the PDE of Equation (6). However, we recommend using an IBM instead. The principal reason for this is that IDEs and PDEs only keep track of the probability distribution of animal locations, but IBMs keep track of the actual location of animals (Wang & Grimm, 2007). This has two advantages. First, if performing numerical simulations, one might as well keep as much realism in them as possible (i.e. why not use an IBM?). Second, animals will respond to the actual (past and present) locations of themselves and other animals, not a distribution that reflects the probability of all possible locations that each animal could have taken. For analytic approaches, PDEs are the best tool and we will discuss this more in Section 4.

Writing code for an IBM depends a lot on the specific situation that is being modelled, especially for interacting objects. We give a basic example in Supplementary Appendix G, of animals that have a mutual avoidance tendency and attraction to a single static resource layer, to help the uninitiated get started. However, we caution the reader that construction of an IBM for their specific situation is likely to require quite significant thought and modification/re-writing of this simple model, which will depend on the particular structure of Equation (1).

### 3.4 Goodness-of-fit analysis from emergent patterns

Having described tools for ascertaining emergent spatial patterns from models parametrised at a finer spatio-temporal scale, we now turn our attention to what we can learn from this process. A principle aim is to examine the extent to which these fine-scale processes capture the observed broad-scale patterns. This process can help reveal missing features from the model and generate new hypotheses (Potts *et al.*, 2022a).

An example of this is shown in Figure 3 using simulated data. In this example, we assume the following movement kernel describes the movement of animal *j*, for *j* ∈ {1, 2, 3, 4}

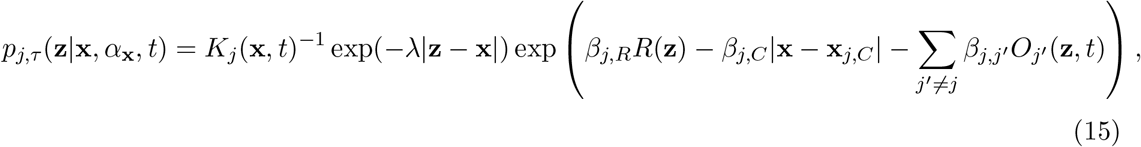

where *R*(**z**) is a resource layer, **x**_*j,C*_ is a localising point for animal *j* and *O_j′_* (**z**, *t*) is the occurence distribution of animal *j*′ at time *t*.

**Fig. 3.**
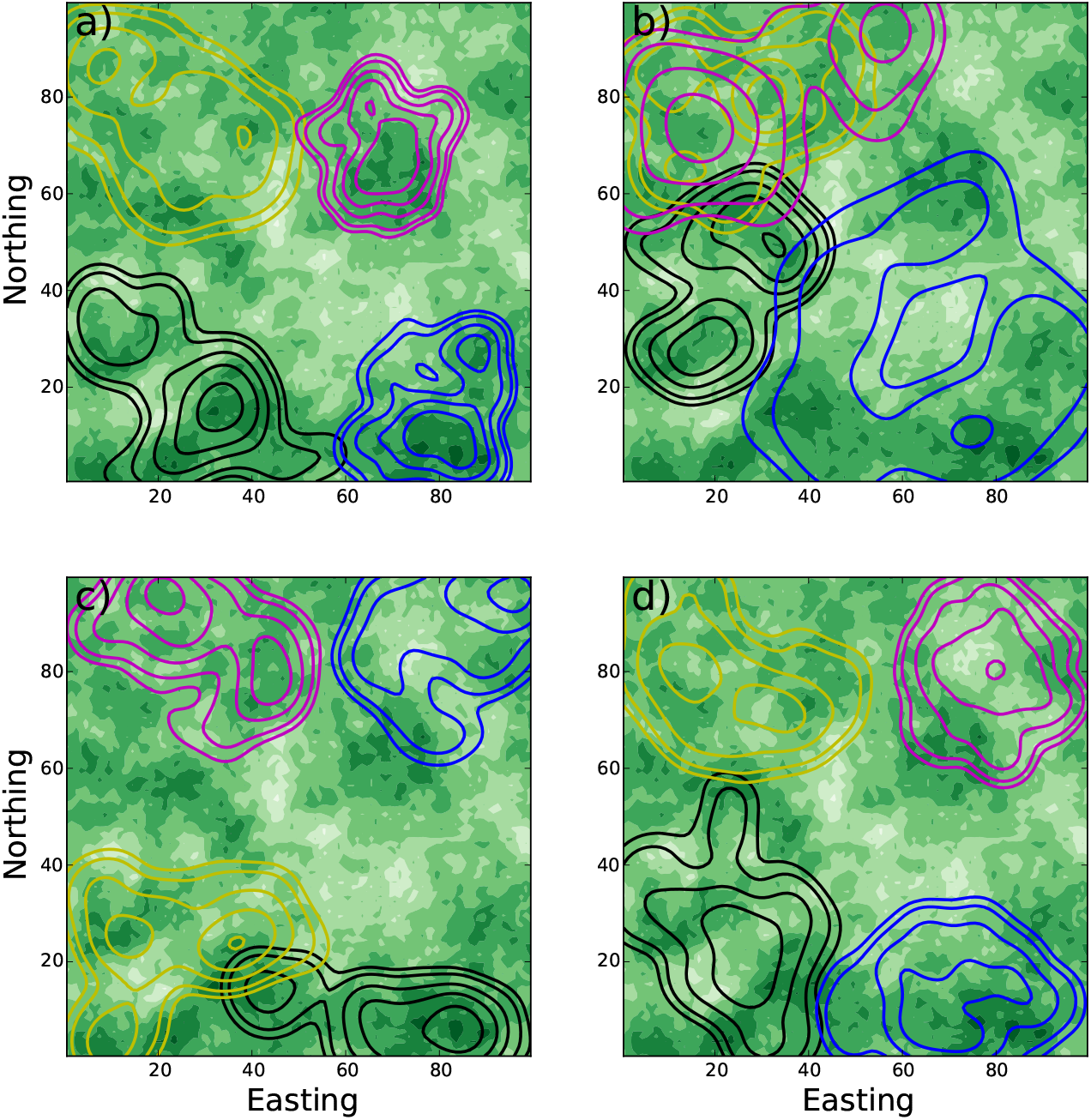
The effect of missing covariates. Empirically-parametrised IBMs can be used to detect missing features from a step selection model. Panel (a) shows some simulated data from four animals in a 100 × 100 box, whose movement is governed by three things: mutual avoidance, attraction to resources, and central place attraction. The green shades represent the resource layer (darker green implies better resources), the contours give UD of animal locations, one colour for each animal (contours at heights 0.0001, 0.0002, 0.0005, 0.001, 0.002 from the outside in). Panel (b) shows the UD when mutual avoidance is removed from the model. Panel (c) has central place attraction removed. Panel (d) does not include resources. If researchers have parametrised an SSF model but missed a key covariate governing space use then they may be able to gain insight into what that missing covariate is by comparing simulated and empirical UDs, in the same way as we might compare Panel (a) with Panels (b-d).

Figure 3a shows the KDE distributions of each simulated animal, moving according to an IBM based on the movement kernel in Equation (15). Figure 3b-d shows the distributions of simulated animals where one of the covariates is missing. There are some technical considerations when constructing an IBM of animals that interact through their occurence distribution (Potts *et al.*, 2022a). In short, one needs to think about how to construct *O_j′_*(**z**, *t*) at each step of the simulation, which involves not just the locations where the animal makes a turn but also locations inbetween turns. One way to deal with this is to simulate a stepping-stone process (Avgar *et al.*, 2013, 2016). Details of how we have constructed such a stepping-stone process from an example system of coupled movement kernels are given in Supplementary Appendix G.

Comparing panels in Figure 3 reveals that a failure to incorporate social interactions (i.e. *β_j,j′_* = 0) leads to more overlap between home ranges than is actually the case (compare panels (a) and (b)), a failure to incorporate localising tendency (i.e. *β_j,C_* = 0) leads to UDs emerging in the wrong place (panels (a) and (c)), and a failure to include the resource layer (i.e. *β_j,R_* = 0) leads to UDs that fail to grasp the environmental heterogeneity in *R*(**z**) (panels (a) and (d)). This is perhaps quite obvious in the omniscient situation of simulated data. However, if in a real situation using empirical data, a researcher has not realised about one of these features, and then has parametrised a model that does not include that feature, then comparing empirical data on space use to emergent patterns from simulations of that model can help reveal this missing feature (Potts *et al.*, 2022a).

As well as visual examination of discrepancies between data and IBM output, various metrics can be calculated to assess goodness of fit. Two possible metrics are to (i) compare the UD or OD sizes between the data and the IBM output, and (ii) to measure the UD or OD overlap between data and IBM. UD size can be measured using any number of metrics, but the locational variance is perhaps the simplest, as it is proportional to standard measures, like 95% KDE, but does not require interpolation or smoothing. Following (Fieberg & Kochanny, 2005), we recommend using Bhattacharyya’s Affinity to measure UD overlap. Details of all these methods are given in Potts *et al.* (2022a).

## 4 Exploring emergent patterns

Whilst IBMs are valuable for comparing model output with data (Section 3.4) they are not always so amenable to mathematical analysis. This is where PDEs come into their own. There is a wealth of techniques for analysing PDEs in the applied mathematics literature (Murray, 2003; Robinson & Pierre, 2003; Buttenschön & Hillen, 2021; Evans, 2022). Many of these are quite technically demanding, and so our best ‘how to’ suggestion for those who do not have a deep background in applied mathematics is to form collaborations with those who do. The trick for forming such collaborations is to know broadly the sort of questions that can be answered by mathematical techniques and how to phrase them in the language of applied maths in a way that might entice collaborators, whilst at the same time keeping firmly grounded in ecological and natural history knowledge. Our philosophy here will be to try to explain how to do this, with the ultimate aim of helping readers form useful collaborations, rather than doing the mathematics themselves.

Perhaps the most elementary technique in pattern formation analysis of PDEs is linear stability analysis (LSA; also sometimes called Turing pattern analysis, after Turing (1952)). This technique asks the following question: if a system is homogeneous (in our case this means the utilisation distribution of each animal/group/population is the whole landscape) and is then perturbed slightly (which will happen naturally as animals move), will those perturbations grow in time? In practice, this means that UDs will segregate or aggregate spontaneously. Therefore it can be used to answer questions such as whether avoidance processes are sufficiently strong to cause territorial segregation, or whether attraction processes are sufficient to enable aggregations to emerge spontaneously. Such analysis may also help researchers to separate-out the effect of social interactions on spatial distribution patterns from environmental interactions.

A second question that can be answered by LSA is: as the patterns grow from a homogeneous state, will they oscillate? This means that any segregations or aggregations that emerge will not be stationary but move around in space. This is of key importance in measuring UDs from data, because if a collection of animals have movement processes that lead to perpetually-fluctuating space use patterns, then this has to be taken into account when measuring UDs from data. For this, one has to consider a set of locations across a time window. The size of this window should be determined by the natural period of any emergent oscillatory patterns.

The next question, which requires tools beyond LSA, is whether patterns are likely to form suddenly as parameters change. This means that a small environmental perturbation might give rise to a dramatic change in the structure of UDs (i.e. a ‘tipping point’). Figure 4b gives an example of this (in the context of IBMs) whereby an increasing tendency for attraction to conspecifics leads to a sudden switch in spatial distribution from homogeneous to highly-aggregated (which could, for example, be driven by increased fear of predation). The existence of these sorts of tipping points can be ascertained by a variety of techniques, perhaps the most well-used of which is *weakly non-linear analysis* (others include *Crandall-Rabinowitz bifurcation theory* and *centre manifold theory* and different mathematicians have different tastes regarding which to use, so it is valuable to be aware of the nomenclature). These techniques can determine whether the point at which patterns start to form (known as the *bifurcation point*) is *supercritical*, meaning that the size of the patterns is continuously dependent on the underlying process (Figure 4a), or *subcritical,* meaning there is a discontinuous switch from no patterns to patterns (Figure 4b).

**Fig. 4.**
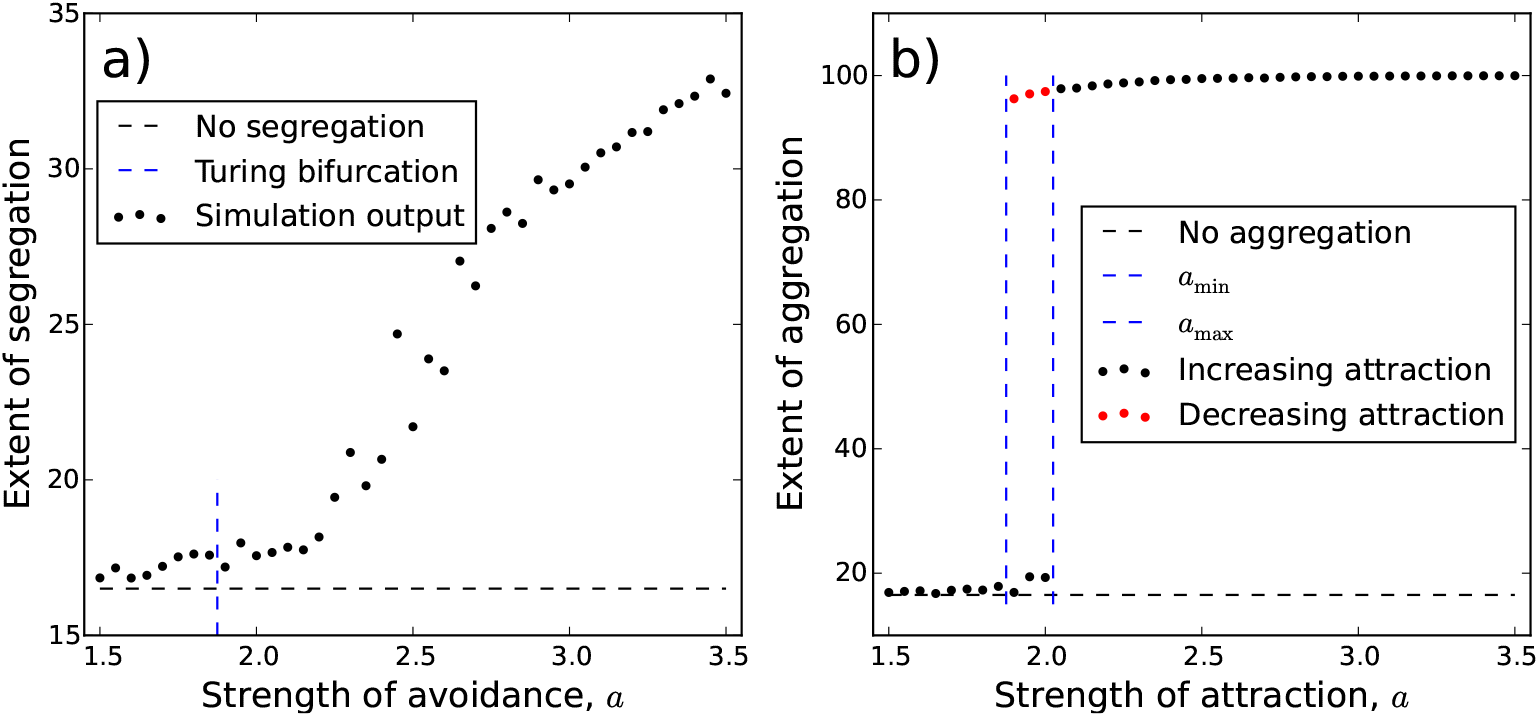
Transition from homogeneity to heterogenous patterns. Panels (a) (resp. Panel (b)) shows simulation output from an stochastic IBM consisting of two mutually-avoiding (resp. mutually-attracting) populations. When the strength of avoidance (resp. attraction) is low, the populations are well-mixed, indistinguishable from non-interacting populations. As this strength is increased past the Turing bifurcation point, the populations begin to segregate (resp. aggregate). Whilst the segregation patterns in Panel (a) emerge in a continuous fashion, the aggregation patterns in Panel (b) appear quite suddenly, and a hysteresis effect is observed. The precise definitions of the quantities in these plots are given in Supplementary Appendix H.

The subcritical case is often accompanied by a hysteresis phenomenon, whereby the existence of patterns depends upon the history of the system. For example, in Figure 4b, in the range *a*_min_ < *a* < *a*_max_, it is possible to see either aggregations or homogeneous UDs depending on the history of the system. A real life example might be of a population of herbivores in the presence of a predation risk that is initially mild, but grows steadily, causing them to increase their tendency to move towards one another for safety (given by the parameter *a*). At some point the predation risk becomes sufficiently high that *a* > *a*_max_ and an aggregation forms. Suppose that after some time the predation risk stops increasing and instead starts to decrease (perhaps due to human intervention or predator disease). Then a decreases. However, when a decreases past *a*_max_, the aggregations do not yet collapse. Indeed, not until *a* < *a*_min_ does this collapse happen and the herbivores return to their original, homogeneously-spread state.

Whilst the mathematical tools of PDE analysis enable rigorous quantification of UD pattern formation properties, the downside is that various approximations are made when moving from the movement kernel of Equation (1) to the PDE of Equation (6). It is therefore valuable to check that the pattern formation properties observed in PDE analysis are also observed in the more realistic case of an IBM. Recent research has begun to develop techniques for doing this, tailored to the specific case of understanding emergent space use patterns from animal movement processes (Potts *et al.*, 2022b). This research shows both how to relate IBMs to PDEs in a rigorous fashion, and gives methods for determining whether patterns emerge, whether they are stationary or fluctuating, aggregative or segregative. Indeed, Figure 4 gives an example of the output of such techniques. In particular, Figure 4a shows the analytically-computed Turing bifurcation point of the PDE system, which is close to where the IBMs bifurcate from homogeneous to heteogeneous patterns.

## 5 Discussion

### 5.1 Why scale up?

We have described various existing techniques for scaling-up from step selection to broader-scale space use patterns, some of which require some relatively involved mathematical and/or numerical analysis. So what is the value in learning and using these techniques? We highlight two key points.

1. **Prediction in systems with feedbacks.** As shown in Sections 2.2 and 3.3, if there are two or more variables that each affect one another, so that there is no *a priori* demarcation into explanatory and response variables, then correlative models alone are insufficient for making predictions (including resource selection analysis (RSA) and many species distribution models (SDMs)). Instead one needs a dynamic model. Step selection provides a technique for parametrising such models and the methods described here provide techniques for analysing their emergent features. An example of this might be predicting the effect of rewilding strategies on ecosystem restoration. For example, domestic herbivores may be allowed to roam more freely, leading to a more heterogeneous vegetation layer, which in turn affects their movements and distributions.
2. **Testing for missing covariates in movement models.** Step selection analysis can demonstrate which of a predetermined set of variables covary with movement. However, it cannot ascertain whether the user has focused on the correct set of variables for describing the animal’s movement. By propogating the resulting movement kernel forwards in time, we can discover the extent to which it can predict longer-term patterns. Any discrepancy between predictions and data can be used to inform further model development, as in Section 3.4, and also where best to concentrate data gathering efforts.

Underlying both of these is a conceptual move from uncovering predictors to building predictive models. Correlative models, such as SSA, RSA, and SDMs, are well-developed for uncovering predictors, but they are less developed regarding making actual predictions. Accurate model predictions require that the underlying models both capture the dynamics correctly (Point 1 above) and contain all the necessary mechanisms required for accurate predictions (Point 2 above). Prediction in spatial animal ecology is notoriously tricky (Hao *et al.*, 2019) and requires significant future development. However, by plugging these two conceptual gaps, the methods described here should provide an important step in improving the predictive power of animal space use models.

### 5.2 Future directions

Building models of animal movement that are able to predict broad-scale space use patterns is a fundamental goal of movement ecology, with a huge range of potential applications to conservation and management (see Introduction). However, there is still a long way to go before accurate predictions from fine-grained movement models become widely possible. We end with a few ideas for where we would like to see these methods going.

1. **Application across taxa and geography.** Many of the methods described here are relatively new and have yet to be applied in earnest to a wide range of datasets. Broad application across different data and research groups is perhaps the best way to ascertain the value of these methods and discover practical ideas for improvement. Ideally, these would include different taxa, populations in contrasting environments, and data sampled over different times scale and intensities.
2. **Biologging data.** Alongside movement, modern datasets often contain a wealth of biologging data, such as heart rate, acceleration, body temperature, and neurological sensors (Williams *et al.*, 2020). These can help inform the behavioural mode of animals, which in turn affects how they move and use space. Methods to incorporate these into movement models, ascertaining the extent to which they feed up to broad space use patterns, is a key research frontier (Klappstein *et al.*, 2021).
3. **Continuous time formulations.** Animals move in continuous time, they may make decisions at any point in time, data may be gathered at completely different points in time, all of which makes a continuous time framework appealing (Parton *et al.*, 2016). In Supplementary Appendix I, we discuss some current efforts to this end, and some possible ways forward. It is also worth mentioning that methods from Sections 3.4 and 4 are not implicitly tied to a discrete time framework, so it would be valuable to examine how to use these in the context of continuous time models.
4. **Mathematical analysis of emergent phenomena.** Our mathematical understanding of emergent phenomena from moving, interacting populations is still in relative infancy (Eftimie, 2018; Potts & Lewis, 2019). Specifically, efforts are required that explicitly tie these into empirically-measured movement processes. This will require strong collaborations between ecologists and applied mathematicians.

## Code availablity

All code can be found on GitHub at https://github.com/jonathan-potts/HowToScaleUp.

## Author contributions

JRP and LB developed the idea for this review. JRP led the writing of the manuscript. Both authors contributed critically to the drafts and gave final approval for publication.

## Acknowledgements

JRP acknowledges support of Engineering and Physical Sciences Research Council (EPSRC) grant EP/V002988/1. We thank Roberto Salguero-Gomez for encouraging us to write this guide and for valuable comments during the process of writing.

