## Supplementary Appendix for "How to scale up from animal movement decisions to spatio-temporal patterns: an approach via step selection"

**2 Department of Biosciences, College of Science, Swansea University, Singleton Park, Swansea, Wales SA2 8PP, UK**

**3 Centre for Biomathematics, College of Science, Swansea University, Singleton Park, Swansea, Wales SA2 8PP, UK**

#### Supplementary Appendix A: Example movement kernels

Here, we give a few foundational examples of movement kernels, to aid readers in understanding the notation used in Section 2.1 of the Main Text and throughout. These will be necessarily simplified models of animal behaviour (i.e. not very realistic), to minimise the mathematical baggage for pedagogical purposes. However, they form the building blocks for more complex and realistic models, which can be captured with the framework we detail in the Main Text.

To start, suppose we have an animal in a completely static environment consisting of a single resource layer, that has a tendency to move towards higher values of that layer (e.g. a herbivore in an environment of heterogeneous vegetation, perhaps measured using Normalised Difference Vegetation Index, NDVI). As well as this tendency, suppose there is some 'random noise' in our movement model (which should be thought of things we are not explicitly modelling, rather than actual random movement, which is probably rare in animals). Suppose we have recorded locations every $\tau$ minutes.

Putting this situation into the notation of Equation (1) of the Main Text, we first notice that there is only one resource layer, $R(\mathbf{x})$, which means that $n = 1$ so the vector $\mathbf{Z}$ only has one entry, $Z_1$. Since the animal has a tendency to move *towards* higher values of the resource layer, $Z_1$ is dependent only on the end-point of the step, given by $\mathbf{z}$. Since the resource layer is assumed to be static, $Z_1$ does not depend upon time. Therefore $Z_1(\mathbf{x}, \mathbf{z}, t) = R(\mathbf{z})$, and this is the value of the resource at location $\mathbf{z}$ (e.g. the NDVI at $\mathbf{z}$). We also write $\beta_R$ instead of $\beta_1$, to emphasise that this value relates to the resource layer, $R$. The 'random noise' can be modelled by assuming that, in the absence of any environmental heterogeneity, the animal moves on average $l$ metres in $\tau$ minutes (i.e. the average 'step length' is $l$), that this length is exponentially distributed, and that the direction is chosen uniformly at random. Then we set $\phi_\tau(\mathbf{z}, \mathbf{x}, \alpha_\mathbf{x}, t) = \exp(-\lambda|\mathbf{z} - \mathbf{x}|)$, where $\lambda = 1/l$. Notice that the step length is $|\mathbf{z} - \mathbf{x}|$, by definition, which is the distance between $\mathbf{z}$ and $\mathbf{x}$. Notice also that this functional form for $\phi_\tau$ is independent of $\alpha_\mathbf{x}$ (the bearing on which the animal arrived at $\mathbf{x}$, see Fig. 1 from the Main Text), as the direction is chosen uniformly at random in the absence of environmental heterogeneity. In conclusion, the movement kernel for this situation, in the form of Equation (1) from the Main Text, is

$$p_\tau(\mathbf{z}|\mathbf{x}) = \frac{\exp(-\lambda|\mathbf{z} - \mathbf{x}|)\exp[\beta_R R(\mathbf{z})]}{\int_\Omega \exp(-\lambda|\mathbf{z} - \mathbf{x}|)\exp[\beta_R R(\mathbf{z})]\mathrm{d}\mathbf{z}}, \tag{S.1}$$

where $p_\tau(\mathbf{z}|\mathbf{x}, \alpha_\mathbf{x}, t) = p_\tau(\mathbf{z}|\mathbf{x})$ as the function in this situation is independent of $\alpha_\mathbf{x}$ and $t$. The denominator here ensures that $p_\tau(\mathbf{z}|\mathbf{x})$ is an actual probability density function, i.e. that it integrates to 1 with respect to $\mathbf{z}$. In practice, if the animal is modelled as moving on a raster layer (with a discrete set of points), one replaces the integral with a sum, to give

$$P_\tau(s|s') = \frac{\exp(-\lambda|s - s'|)\exp[\beta_R R(s)]}{\sum_{s \in S} \exp(-\lambda|s - s'|)\exp[\beta_R R(s)]\mathrm{d}s}, \tag{S.2}$$

which is the probability of moving from $s'$ to $s$ in a timestep of length $\tau$, and $S$ is the set of pixels in the raster layer. Code is attached in `htsu_sim_path_ex1.py` (in `Python`) and `htsu_sim_path_ex1.R` (in `R`) for simulating Equation (S.2) on an example layer given by `random_field_100.inp`. This layer is the one displayed in Figs. 1-3 in the Main Text. All code is also on GitHub at https://github.com/jonathan-potts/HowToScaleUp.

To add another layer of complexity, suppose that the animal has, in addition to the above assumptions, some autocorrelation in movement. For example, suppose that, in the absence of environmental heterogeneity, the animal is more likely to choose a direction that is similar to their existing direction of movement than one that is dissimilar. If we denote the bearing from $\mathbf{x}$ to $\mathbf{z}$ by $\alpha_{(\mathbf{x},\mathbf{z})}$, then we set $\phi_\tau(\mathbf{z},\mathbf{x},\alpha_\mathbf{x},t) = \exp(-\lambda|\mathbf{z}-\mathbf{x}|)\exp[\kappa\cos(\alpha_\mathbf{x}-\alpha_{(\mathbf{x},\mathbf{z})})]$, which can also be written as $\exp[-\lambda|\mathbf{z}-\mathbf{x}|+\kappa\cos(\alpha_\mathbf{x}-\alpha_{(\mathbf{x},\mathbf{z})})]$. The reason for this is that $\cos(\alpha_\mathbf{x}-\alpha_{(\mathbf{x},\mathbf{z})})$ is maximised when $\alpha_\mathbf{x}=\alpha_{(\mathbf{x},\mathbf{z})}$ (i.e. the directions of motion prior to arriving at $\mathbf{x}$ is the same as the direction from $\mathbf{x}$ to $\mathbf{z}$) and minimised when $\alpha_\mathbf{x}$ and $\alpha_{(\mathbf{x},\mathbf{z})}$ differ by $\pi$ radians (180°). The parameter $\kappa$ denotes the strength of the bias towards carrying on moving in a similar direction. This formalism is identical to the von Mises distribution, oft-used for modelling correlated random walks (Turchin, 1998; Codling *et al.*, 2008). The movement kernel in this case is [Equation (1) from the Main Text]

$$p_\tau(\mathbf{z}|\mathbf{x}) = \frac{\exp[-\lambda|\mathbf{z}-\mathbf{x}|+\kappa\cos(\alpha_\mathbf{x}-\alpha_{(\mathbf{x},\mathbf{z})})]\exp[\beta_R R(\mathbf{z})]}{\int_\Omega \exp[-\lambda|\mathbf{z}-\mathbf{x}|+\kappa\cos(\alpha_\mathbf{x}-\alpha_{(\mathbf{x},\mathbf{z})})]\exp[\beta_R R(\mathbf{z})]\mathrm{d}\mathbf{z}}. \tag{S.3}$$

Again, in a raster environment, where we model the animal as moving from pixel to pixel on a lattice $S$, we can write

$$P_\tau(s|s') = \frac{\exp[-\lambda|s-s'|+\kappa\cos(\alpha_{s'}-\alpha_{(s',s)})]\exp[\beta_R R(s)]}{\sum_{s\in S}\exp[-\lambda|s-s'|+\kappa\cos(\alpha_{s'}-\alpha_{(s',s)})]\exp[\beta_R R(s)]\mathrm{d}s}. \tag{S.4}$$

Code for simulating this movement kernel is given in `htsu_sim_path_ex2.py` (in `Python`) and `htsu_sim_path_ex2.R` (in `R`).

For our third and final example, suppose we have two animals, each moving as in the previous example, but with the additional aspect of having a tendency to avoid recent locations of each other, given by the occurence distrubution over the past $T$ units of time. We assume that each animal advertises their recent locations through marking the environment (e.g. using urine or faeces). Then, in addition to $Z_1$, denoting the static environmental layer, we need layers $Z_{2,1}$ and $Z_{2,2}$ denoting the occurence distribution of animal 2 as percieved by animal 1, and the occurence distribution of animal 1 as percieved by animal 2, respectively. These change over time, so depend upon $t$, and affect the probability of choosing a location at the end of a step, so depend upon $\mathbf{z}$. Thus we write $Z_{2,1} = Z_{2,1}(\mathbf{z}, t)$ and $Z_{2,2} = Z_{2,2}(\mathbf{z}, t)$.

Since there are two animals, we also need to decorate the symbols $p_\tau$ and $\phi_\tau$ with an additional subscript denoting the number of the animal, as in Equation (3) of the Main Text. Hence $p_\tau$ (resp. $\phi_\tau$) becomes $p_{1,\tau}$ (resp. $\phi_{1,\tau}$) for Animal 1 and $p_{2,\tau}$ (resp. $\phi_{2,\tau}$) for Animal 2. In summary the movement kernels become (Equation (3) from the Main Text)

$$p_{\tau,1}(\mathbf{z}|\mathbf{x}, t) = \frac{\exp[-\lambda|\mathbf{z}-\mathbf{x}| + \kappa\cos(\alpha_{\mathbf{x}} - \alpha_{(\mathbf{x},\mathbf{z})})]\exp[\beta_R R(\mathbf{z}) + \beta_2 Z_{2,1}(\mathbf{z}, t)]}{\int_\Omega \exp[-\lambda|\mathbf{z}-\mathbf{x}| + \kappa\cos(\alpha_{\mathbf{x}} - \alpha_{(\mathbf{x},\mathbf{z})})]\exp[\beta_R R(\mathbf{z}) + \beta_2 Z_{2,1}(\mathbf{z}, t)]\mathrm{d}\mathbf{z}} \tag{S.5}$$

$$p_{\tau,2}(\mathbf{z}|\mathbf{x}, t) = \frac{\exp[-\lambda|\mathbf{z}-\mathbf{x}| + \kappa\cos(\alpha_{\mathbf{x}} - \alpha_{(\mathbf{x},\mathbf{z})})]\exp[\beta_R R(\mathbf{z}) + \beta_2 Z_{2,2}(\mathbf{z}, t)]}{\int_\Omega \exp[-\lambda|\mathbf{z}-\mathbf{x}| + \kappa\cos(\alpha_{\mathbf{x}} - \alpha_{(\mathbf{x},\mathbf{z})})]\exp[\beta_R R(\mathbf{z}) + \beta_2 Z_{2,2}(\mathbf{z}, t)]\mathrm{d}\mathbf{z}}, \tag{S.6}$$

in continuous space or

$$p_{\tau,1}(\mathbf{z}|\mathbf{x}, t) = \frac{\exp[-\lambda|\mathbf{z}-\mathbf{x}| + \cos(\alpha_{\mathbf{x}} - \alpha_{(\mathbf{x},\mathbf{z})})]\exp[\beta_R R(\mathbf{z}) + \beta_2 Z_{2,1}(\mathbf{z}, t)]}{\sum_{\mathbf{z}\in S} \exp[-\lambda|\mathbf{z}-\mathbf{x}| + \cos(\alpha_{\mathbf{x}} - \alpha_{(\mathbf{x},\mathbf{z})})]\exp[\beta_R R(\mathbf{z}) + \beta_2 Z_{2,1}(\mathbf{z}, t)]\mathrm{d}\mathbf{z}}. \tag{S.7}$$

$$p_{\tau,2}(\mathbf{z}|\mathbf{x}) = \frac{\exp[-\lambda|\mathbf{z}-\mathbf{x}| + \cos(\alpha_{\mathbf{x}} - \alpha_{(\mathbf{x},\mathbf{z})})]\exp[\beta_R R(\mathbf{z}) + \beta_2 Z_{2,2}(\mathbf{z}, t)]}{\sum_{\mathbf{z}\in S} \exp[-\lambda|\mathbf{z}-\mathbf{x}| + \cos(\alpha_{\mathbf{x}} - \alpha_{(\mathbf{x},\mathbf{z})})]\exp[\beta_R R(\mathbf{z}) + \beta_2 Z_{2,2}(\mathbf{z}, t)]\mathrm{d}\mathbf{z}}, \tag{S.8}$$

in discrete space. Simulating these engenders technical difficulties regarding how to construct the occurence distribution over time. These difficulties are resolved in Potts *et al.* (2022a) and we explain how to construct an IBM from Equations (S.7-S.8) in Supplementary Appendix G.

These three examples are intended to give readers a road into understanding the general formalisms from Equations (1) and (3) of the Main Text. We end by explaining how to relate Equation (1) from the Main Text to Equation (13) from Fieberg *et al.* (2021), to enable readers to interface the two 'How to' articles. Equation (13) from Fieberg *et al.* (2021) reads as (after correcting a typographical error in the denominator)

$$u(s, t+\Delta t)|u(s', t) = \frac{w(X(s); \beta(\Delta t))\phi(s, s'; \gamma(\Delta t))}{\int_G w(X(\tilde{s}); \beta(\Delta t))\phi(\tilde{s}, s'; \gamma(\Delta t))d\tilde{s}}. \tag{S.9}$$

To relate this to Equation (1) from the Main Text requires the following transformations

$$s \mapsto \mathbf{z}, \qquad s' \mapsto \mathbf{x}, \qquad \Delta t \mapsto \tau, \qquad X(s) \mapsto \mathbf{Z}(\mathbf{x}, \mathbf{z}, t), \qquad \beta(\Delta t) \mapsto \boldsymbol{\beta},$$
$$\phi(s, s'; \gamma(\Delta t)) \mapsto \phi_\tau(\mathbf{z}, \mathbf{x}, \alpha_\mathbf{x}, t), \qquad u(s, t + \Delta t)|u(s', t) \mapsto p_\tau(\mathbf{z}|\mathbf{x}, \alpha_\mathbf{x}, t).$$

A few things to note:

– In replacing $X(s)$ by $\mathbf{Z}(\mathbf{x}, \mathbf{z}, t)$, we allow for dependence of any landscape feature on the start of the step, the end of the step, and time, rather than just the end of the step.

– By using the notation $\beta(\Delta t)$, Fieberg *et al.* (2021) emphasise the fact that $\beta$ may be different depending on the step time ($\tau$ in our work, $\Delta t$ in theirs). Whilst this is certainly true, and should be borne in mind if $\tau$ were changed, we have instead fixed $\tau$ in our explanations, so do not explicitly include this dependency in our notation.

– Our function $\phi_\tau$ is explicitly dependent on the previous direction of travel, $\alpha_\mathbf{x}$. This is discussed in Fieberg *et al.* (2021), but is not explicit in their notation.

– Likewise, our function $p_\tau$ is explicitly dependent on the previous direction of travel, $\alpha_\mathbf{x}$. Again, this appears to be implicit in Fieberg *et al.* (2021), but not made explicit in their notation.

### Supplementary Appendix B: Alterations required for high frequency data

Recall the movement kernel from Equation (1) of the Main Text:

$$p_\tau(\mathbf{z}|\mathbf{x},\alpha_\mathbf{x},t) = \frac{\phi_\tau(\mathbf{z},\mathbf{x},\alpha_\mathbf{x},t)\exp[\boldsymbol{\beta}_\tau \cdot \mathbf{Z}(\mathbf{x},\mathbf{z},t)]}{\int_\Omega \phi_\tau(\mathbf{z},\mathbf{x},\alpha_\mathbf{x},t)\exp[\boldsymbol{\beta}_\tau \cdot \mathbf{Z}(\mathbf{x},\mathbf{z},t)]\mathrm{d}\mathbf{z}}. \tag{S.10}$$

In many SSA studies, the movement kernel is parametrised by using pairs of consecutively-measured locations for $\mathbf{x}$ and $\mathbf{z}$, with perhaps the prime example being GPS locations. However, there are two problems with this. First, animals will almost certainly not be making their decisions about movement at the precise moments the tracker (GPS or otherwise) measures their location. Second, increasingly many modern datasets are of such high frequency that it is completely unreasonable to suggest that the animal is making discrete movement decisions at anything like the frequency of the data. Indeed, many modern datasets have locations measured multiple times per second. Most of these locations will simply reflect the animal continuing to move along the trajectory on which it decided to move many time steps before.

Therefore, if one is to use such high frequency data for step selection, it is important to rarefy the data first. Following Munden *et al.* (2021), we suggest that the best way to rarefy a high frequency movement path for use with SSA is to detemine the places where the animal turned, as these are more likely to correspond to movement decisions than any other measured location (Wilson *et al.*, 2013). A rapid algorithm for turning-point identification is given in Potts *et al.* (2018), the supporting information of which contains code for running the algorithm, in `R` and `Python`, as well as instructions and a simple case study.

Performing SSA on a time series of inferred turning points requires some modification of the usual SSA technique, since consecutive turning points will typically not be separated by a constant time interval, $\tau$. In practice this means that the movement kernel from Equation (1) in the Main Text needs to be replaced with the following time-varying modification

$$p(\mathbf{z}, \tau | \mathbf{x}, \alpha_{\mathbf{x}}, t) = \frac{\phi(\mathbf{z}, \mathbf{x}, \alpha_{\mathbf{x}}, t, \tau) \exp[\boldsymbol{\beta} \cdot \mathbf{Z}(\mathbf{x}, \mathbf{z}, t)]}{\int_{\Omega} \left( \int_0^{\infty} \phi(\mathbf{z}, \mathbf{x}, \alpha_{\mathbf{x}}, t, \tau) \exp[\boldsymbol{\beta} \cdot \mathbf{Z}(\mathbf{x}, \mathbf{z}, t)] \mathrm{d}\tau \right) \mathrm{d}\mathbf{z}}. \tag{S.11}$$

This can be parametrised using an extension of SSA called 'time-varying integrated SSA' (tiSSA) and details of this procedure are given in Munden *et al.* (2021).

#### Supplementary Appendix C: Alterations required for correlated movement

Recall the master equation given in Equation (4) of the Main Text:

$$u(\mathbf{z}, t+\tau) = \int_{\Omega} p_{\tau}(\mathbf{z}|\mathbf{x}, t) u(\mathbf{x}, t) \mathrm{d}\mathbf{x}. \tag{S.12}$$

Here, the assumption is that $p_\tau$ is independent of $\alpha_{\mathbf{x}}$ (i.e. no correlation in movement), so that $p_\tau(\mathbf{z}|\mathbf{x}, \alpha_{\mathbf{x}}) = p_\tau(\mathbf{z}|\mathbf{x}, t)$.

To incorporate the aspect of correlated movement (i.e the dependence on $\alpha_{\mathbf{x}}$), we need to define the quantity $u_\alpha(\mathbf{x}, \alpha_{\mathbf{x}}, t)$, which is the UD of locations given that the animal has arrived at that location on a bearing of $\alpha_{\mathbf{x}}$. From an ecological perspective, $u_\alpha$ is perhaps a bit of a strange and counter-intuitive quantity, but it makes sense mathematically. Most importantly, we can write down the master equation for $u_\alpha$ as follows

$$u_\alpha(\mathbf{z}, \alpha_{\mathbf{z}}, t+\tau) = \int_0^{2\pi} \left( \int_{\Omega} p_\tau(\mathbf{z}|\mathbf{x}, \alpha_{\mathbf{x}}, t) u_\alpha(\mathbf{x}, \alpha_{\mathbf{x}}, t) \mathrm{d}\mathbf{x} \right) \mathrm{d}\alpha_{\mathbf{x}}, \tag{S.13}$$

and then use this to calculate the UD itself, as

$$u(\mathbf{x}, t) = \int_0^{2\pi} u_\alpha(\mathbf{x}, \alpha_{\mathbf{x}}, t) \mathrm{d}\alpha_{\mathbf{x}}. \tag{S.14}$$

Immediately, we observe that Equations (S.12) and (S.13) are not readily extensible for use with Equation (S.11) instead of Equation (S.10), at least without some careful thought. This is because Equations (S.12) and (S.13) are predicated on having a pre-determined fixed time-interval, $\tau$ between locations. A naïve approach might be to calculate the average value of $\tau$ from data and use that in Equation (S.12). However, we caution that this is a hitherto untested approach (as far as we are aware), and more research needs to be done to ascertain whether it may bias results. We encourage researchers to think about this as yet unsolved problem.

The discrete time formulation, if correlations are included, is constructed as follows. Write the probability of moving from $s'$ to $s$, given that the animal arrives at $s'$ on a bearing of $\alpha_{s'}$, as $P_\tau(s|s', \alpha_{s'}, t)$. Let $U_\alpha(s, \alpha_s, t)$ the probability of an animal being at $s$ at time $t$, having arrived there on a bearing of $\alpha_s$. Then the discrete-space master equation is

$$U_\alpha(s, \alpha_s, t+\tau) = \sum_{i=1}^{M} \sum_{s' \in S} P_\tau \left( s \middle| s', \frac{i}{2\pi}, t \right) U_\alpha \left( s, \frac{i}{2\pi}, t \right), \tag{S.15}$$

which we can compare with Equation (5) from the Main Text.

#### Supplementary Appendix D: Constructing the master equation

Here, we given an example of calculating Equation (5) in the Main Text for a particular movement kernel. For this, we will use the most elementary example from Equation (S.2), whereby the animal is biased towards higher quality resources, described by a single landscape layer. The discrete-space master equation in this case is as follows

$$U(s,t+\tau) = \sum_{s' \in S} \frac{\exp(-\lambda|s-s'|)\exp[\beta_R R(s)]}{\sum_{s \in S} \exp(-\lambda|s-s'|)\exp[\beta_R R(s)]\mathrm{d}s} U(s',t). \tag{S.16}$$

The landscape layer, describing the function $R(s)$, is given by the file `random_field_50.inp`. Code for solving Equation (S.16) over 100 timesteps is given in `htsu_me_ex1.py` (in `Python`) and `htsu_me_ex1.R` (in `R`). For this, we require setting initial conditions, i.e. values of $U(s,0)$. In our code, we set $U(s,0) = 0$ except for the central point of the lattice, $s = (25,25)$, where $U(s,0) = 1$. This models a situation where we know that the animal is in the centre of the landscape at time $t = 0$.

#### Supplementary Appendix E: Calculating the steady state UD using matrix inversion

Here we explain a way in which one can calculate the steady state UD given in Equation (10) of the Main Text, which is the following Equation

$$U_*(s) = \sum_{s' \in S} P_\tau(s|s')U_*(s'). \tag{S.17}$$

This technique has, to our knowledge, never been put into practice in the context of step selection functions, so we do not give detailed instructions for doing this, but it is worth a mention.

First, we need to write Equation (S.17) as a matrix equation. We write $S = \{s_1, \ldots, s_M\}$, where $M$ is the number of elements in $S$ (recall that $S$ is a lattice denoting the study area). The choice of ordering for the lattice points $s_i$ can be picked arbitrarily. Then Equation (S.17) becomes

$$U_*(s_i) = \sum_{j=1}^{M} P_\tau(s_i|s_j)U_*(s_j). \tag{S.18}$$

Writing $\mathbf{U}_* = (U_*(s_1), \ldots, U_*(s_M))^T$ (where the superscript $T$ is for 'transpose'), and defining a matrix $A$ such that the $i,j$-th entry of $A$ is $A_{ij} = P_\tau(s_i|s_j)$, leads to the following matrix equation

$$\mathbf{U}_* = A\mathbf{U}_*, \tag{S.19}$$

where $A$ is independent of $\mathbf{U}_*$. Consequently, $\mathbf{U}_*$ is the eigenvector of $A$ that has eigenvalue 1. It is therefore possible to calculate this steady-state using an eigenvector solver. Whilst this is a relatively trivial observation, we do not know any studies that have applied this method to inferring steady state UDs from SSFs. It is an exact method, so has some appeal, but we suspect that it would typically be computationally demanding in reaslistic situations. However, there is a large and growing body of work on efficient computation of eigenvalue problems (Saad, 2011), so there may yet be value in exploring this approach.

### Supplementary Appendix F: Calculating the steady state UD using the Barnett-Moorcroft method

Recall Equation (11) from the Main Text, which gives the following expression for the steady state utilisation distribution (UD)

$$u_*(\mathbf{x}) = \frac{\exp(\boldsymbol{\beta} \cdot \tilde{\mathbf{Z}}(\mathbf{x})) \int_\Omega \exp(\boldsymbol{\beta} \cdot \tilde{\mathbf{Z}}(\mathbf{z}))\psi_\tau(|\mathbf{x} - \mathbf{z}|)\mathrm{d}\mathbf{z}}{\int_\Omega \left[\exp(\boldsymbol{\beta} \cdot \tilde{\mathbf{Z}}(\mathbf{x})) \int_\Omega \exp(\boldsymbol{\beta} \cdot \tilde{\mathbf{Z}}(\mathbf{z}))\psi_\tau(|\mathbf{x} - \mathbf{z}|)\mathrm{d}\mathbf{z}\right] \mathrm{d}\mathbf{x}}. \tag{S.20}$$

For the movement kernel from Equation (S.1), modelling an animal with a bias to higher values of $R(\mathbf{x})$, the steady state UD is

$$u_*(\mathbf{x}) = \frac{\exp(\beta_R R(\mathbf{x})) \int_\Omega \exp(-\lambda|\mathbf{z} - \mathbf{x}|) \exp(\beta_R R(\mathbf{z}))\mathrm{d}\mathbf{z}}{\int_\Omega \left[\exp(\beta_R R(\mathbf{x})) \int_\Omega \exp(-\lambda|\mathbf{z} - \mathbf{x}|) \exp(\beta_R R(\mathbf{z}))\mathrm{d}\mathbf{z}\right] \mathrm{d}\mathbf{x}}. \tag{S.21}$$

Code for calculating Equation (S.21) is given in `htsu_bm_ud_ex1.py` (in `Python`) and `htsu_bm_ud_ex1.R` (in R). In this code, we approximate the integrals as sums, as explained in Section 3.1 of the Main Text. This code also includes calculations of the estimated UDs from Equations (12) and (13) in the Main Text, but using the movement kernel from Equation (S.1).

In addition to this, the file `htsu_bm_from_ssa.R` (in `R`) both parametrises a movement kernel of the form of Equation (S.1) and then uses that parametrised movement kernel and to predict the steady state UD via Equation (S.21). This demonstrates a complete workflow from step selection analysis through to UD prediction.

As another example, suppose we model an animal with a bias towards higher values of the same resource gradient, and also bias towards a fixed point (e.g. a den or nest site). The movement kernel for this is given in Equation (14) of the Main Text and, by placing this into

Equation (S.20), we return the following expression for the UD

$$u_*(\mathbf{x}) = \frac{\exp(\beta_R R(\mathbf{x}) - \beta_C|\mathbf{x}-\mathbf{x}_C|)\int_\Omega \exp(-\lambda|\mathbf{z}-\mathbf{x}|)\exp(\beta_R R(\mathbf{z}) - \beta_C|\mathbf{z}-\mathbf{x}_C|)\mathrm{d}\mathbf{z}}{\int_\Omega \left[\exp(\beta_R R(\mathbf{x}) - \beta_C|\mathbf{x}-\mathbf{x}_C|)\int_\Omega \exp(-\lambda|\mathbf{z}-\mathbf{x}|)\exp(\beta_R R(\mathbf{z}) - \beta_C|\mathbf{z}-\mathbf{x}_C|)\mathrm{d}\mathbf{z}\right]\mathrm{d}\mathbf{x}}. \tag{S.22}$$

Code for calculating Equation (S.22) is given in `htsu_bm_ud_ex2.py` and `htsu_bm_ud_ex2.R`. This code also includes calculations of the estimated UDs from Equations (12) and (13) in the Main Text, but using the movement kernel from Equation (14) in the Main Text.

We end this appendix with a derivation of Equations (12) and (13) from the Main Text. These are relatively technical compared to the rest of the SI, and it is not necessary to understand these derivations for the purposes of using the techniques described here. However, we include them for completeness, and for the more technical reader. Starting with the derivation of Equation (12), here we are assuming that $\psi_\tau$ is a uniform distribution across $\Omega$. In other words, $\psi_\tau(l) = 1/|\Omega|$, where $|\Omega|$ is the area of $\Omega$. Plugging this into Equation (S.20) gives

$$u_*(\mathbf{x}) = \frac{\exp(\boldsymbol{\beta}\cdot\tilde{\mathbf{Z}}(\mathbf{x}))\frac{1}{|\Omega|}\int_\Omega \exp(\boldsymbol{\beta}\cdot\tilde{\mathbf{Z}}(\mathbf{z}))\mathrm{d}\mathbf{z}}{\int_\Omega\left[\exp(\boldsymbol{\beta}\cdot\tilde{\mathbf{Z}}(\mathbf{x}))\frac{1}{|\Omega|}\int_\Omega \exp(\boldsymbol{\beta}\cdot\tilde{\mathbf{Z}}(\mathbf{z}))\mathrm{d}\mathbf{z}\right]\mathrm{d}\mathbf{x}} = \frac{\exp(2\boldsymbol{\beta}\cdot\tilde{\mathbf{Z}}(\mathbf{x}))}{\int_\Omega \exp(2\boldsymbol{\beta}\cdot\tilde{\mathbf{Z}}(\mathbf{x}))\mathrm{d}\mathbf{x}}, \tag{S.23}$$

which is Equation (12) from the Main Text.

For Equation (13), we assume that $\psi_\tau$ is arbitrarily narrow, which we write as a Dirac-$\delta$ function, $\psi_\tau(l) = \delta(l)$. Then

$$u_*(\mathbf{x}) = \frac{\exp(\boldsymbol{\beta}\cdot\tilde{\mathbf{Z}}(\mathbf{x}))\int_\Omega \exp(\boldsymbol{\beta}\cdot\tilde{\mathbf{Z}}(\mathbf{z}))\delta(|\mathbf{x}-\mathbf{z}|)\mathrm{d}\mathbf{z}}{\int_\Omega\left[\exp(\boldsymbol{\beta}\cdot\tilde{\mathbf{Z}}(\mathbf{x}))\int_\Omega \exp(\boldsymbol{\beta}\cdot\tilde{\mathbf{Z}}(\mathbf{z}))\delta(|\mathbf{x}-\mathbf{z}|)\mathrm{d}\mathbf{z}\right]\mathrm{d}\mathbf{x}} = \frac{\exp(\boldsymbol{\beta}\cdot\tilde{\mathbf{Z}}(\mathbf{x}))}{\int_\Omega \exp(\boldsymbol{\beta}\cdot\tilde{\mathbf{Z}}(\mathbf{x}))\mathrm{d}\mathbf{x}}, \tag{S.24}$$

which is Equation (13) from the Main Text.

### Supplementary Appendix G: Interacting stochastic IBM example

Here we describe how to construct an IBM for animals who interact through responding to one another's recent locations (which in reality could be advertised through marking the terrain, for example). For this, it is valuable to use a nearest neighbour random walk, also sometimes called a 'stepping stone process' (Avgar *et al.*, 2016). In our example, each animal moves on a rectangular lattice and at each step the animal can only move to an adjacent lattice point (left, right, up, or down). Potts *et al.* (2022a) show how to go from a movement kernel (of the type in Equation (1) of the Main Text) to a stepping stone process. At the end of this Appendix, we will show how to do this in our particular example. First, though, we give details of an example IBM.

Suppose each animal has a tendency to move away from the occurrence distribution (OD) of the other and is also biased towards better resources, given by a resource layer $R(s)$, for $s \in S$ where $S$ is a lattice. Then the probabilities, $Q_1(s|s', \Delta t)$ and $Q_2(s|s', \Delta t)$, of moving from one square on the lattice to another in a time step $\Delta t$, for animals 1 and 2 respectively, are written as

$$Q_1(s|s', \Delta t) = \begin{cases} \frac{d}{K_1} \exp\left(\beta_{1,R} R(s) - \beta_{1,2} O_2(s,t)\right), & \text{if } s' \text{ is adjacent to } s, \\ \frac{1-4d}{K_1}, & \text{if } s = s', \\ 0, & \text{otherwise}, \end{cases} \tag{S.25}$$

$$Q_2(s|s', \Delta t) = \begin{cases} \frac{d}{K_2} \exp\left(\beta_{2,R} R(s) - \beta_{2,1} O_1(s,t)\right), & \text{if } s' \text{ is adjacent to } s, \\ \frac{1-4d}{K_2}, & \text{if } s = s', \\ 0, & \text{otherwise}, \end{cases} \tag{S.26}$$

where $0 < d < 0.25$, and $K_1, K_2$ are normalising kernels ensuring the probabilities $Q_1$ and $Q_2$ sum to 1, and the other notation is taken from the Main Text. Code for simulating this is in `htsu_sim_2indivs.py` (in `Python`) and `htsu_sim_2indivs.R` (in `R`) with $d = 0.25$.

Equations (S.25-S.26) describe a stepping stone process that is in a different form from the movement kernel that arises from step selection, as given in Equation (1) of the Main Text. However, they can be formally related. Here, we show how to make this relation for our specific example. The reader is referred to Potts *et al.* (2022a) for the general case.

We begin with the following movement kernels for animals 1 and 2 respectively (cf. Equation (3) from the Main Text)

$$p_{1,\tau}(\mathbf{z}|\mathbf{x}) = [K_1(\mathbf{x}, t)] \exp(-\lambda|\mathbf{z} - \mathbf{x}|) \exp[\beta_{1,R} R(\mathbf{z}) - \beta_{1,2} O_2(s, t)], \tag{S.27}$$

$$p_{2,\tau}(\mathbf{z}|\mathbf{x}) = [K_2(\mathbf{x}, t)] \exp(-\lambda|\mathbf{z} - \mathbf{x}|) \exp[\beta_{2,R} R(\mathbf{z}) - \beta_{2,1} O_1(s, t)]. \tag{S.28}$$

Then the stepping stone process that has the same movement statistics (i.e. the same mean drift and diffusion) is the one given in Equations (S.25-S.26) where

$$d = \frac{\Delta t \int_\Omega |\mathbf{x}|^2 \exp(-\lambda|\mathbf{x}|)\mathrm{d}\mathbf{x}}{\tau(\Delta x)^2 \int_\Omega \exp(-\lambda|\mathbf{x}|)\mathrm{d}\mathbf{x}}, \tag{S.29}$$

for $i = 1, 2$, where $\Delta x$ is the lattice spacing. Notice that there are two time intervals here: $\Delta t$ for the stepping stone process and $\tau$ for the movement kernel. Now, $\tau$ is determined by the data, but we are free to choose $\Delta t$. Indeed, the user needs to choose $\Delta t$ small enough so that $d < 0.25$, otherwise Equations (S.25-S.26) do not give a well-defined probability distribution.

### Supplementary Appendix H: Details behind Figure 4 from the Main Text

The individual based models (IBMs) for Figure 4 from the Main Text follow the structure detailed in Potts *et al.* (2022b). For the purposes of having notation in one place, we recall Equations (1-3) from that paper, which describe the IBM. However, we refer readers to that paper for justification of the model setup, a discussion of the meaning of the various parameters, and details of how to run and analyse these IBMs. Here we just explain how to relate Figure 4 to the precise concepts and notation used in Potts *et al.* (2022b).

Equation (1) from Potts *et al.* (2022b) gives the movement kernel for an individual from population $i$ as follows

$$f(\mathbf{x}, t+\tau|\mathbf{x}', t) = \begin{cases} K_{\mathbf{x}'} \exp\left[\sum_{j=1}^{N} a_{ij}\bar{m}_j^{\delta}(\mathbf{x}, t)\right], & \text{if } |\mathbf{x}-\mathbf{x}'| = l, \\ 0, & \text{otherwise,} \end{cases} \tag{S.30}$$

where $K_{\mathbf{x}'}$ is a normalising constant. Here, $m_j^{\delta}$ is the density of marks left on the terrain by the individual, which are deposited and decay according to the following equation (Equation (3) from Potts *et al.* (2022b))

$$m_i(\mathbf{x}, t+\tau) = (1-\mu_\tau)m_i(\mathbf{x}, t) + \rho_\tau \mathcal{N}_i(\mathbf{x}, t), \tag{S.31}$$

Then $\bar{m}_j^{\delta}$ us the average mark density, given by

$$\bar{m}_j^{\delta}(\mathbf{x}, t) = \frac{1}{|S_\delta|}\sum_{\mathbf{z}\in S_\delta} m_j(\mathbf{x}+\mathbf{z}, t), \tag{S.32}$$

where $S_\delta = \{\mathbf{z} \in \Lambda : |\mathbf{z}| < \delta\}$ is the set of lattice sites that are within a distance of $\delta$ from 0 and $|S_\delta|$ is the number of lattice sites in $S_\delta$.

The IBMs used to construct Figure 4 have the following parameters set as constant: $\rho_\tau =$

0.01, $\mu_\tau = 0.06$, and $\delta = 5$ lattice sites. We also assume that there are two populations, and $a_{12} = a_{21}$, $a_{11} = a_{22} = 0$. Simulations take place on a $25 \times 25$ square lattice with periodic boundary conditions and there are 100 animals in each population. The 'strength of avoidance, $a$', referenced in Figure 4a, is $a = -a_{12}$ and the 'strength of attraction, $a$', referenced in Figure 4b, is $a = a_{12}$. The 'extent of segregation' and 'extent of aggregation' are the values $A^*_{1,5}$ calculated in Potts *et al.* (2022b) (see in particular Figure 3 from that paper). The value of 'no aggregation' and 'no segregation' is the value $A_{rw}$ from Potts *et al.* (2022b), which gives the value of $A^*_{1,5}$ obtained from a system of individuals moving as independent random walkers.

Each black dot in Figure 4 was calculated in an identical way to the corresponding dots in Potts *et al.* (2022b, Figure 3), by beginning with $a = 1.5$, running the simulation until its dynamics are 'stable' (in the sense defined precisely in Potts *et al.* (2022b)), then increasing $a$ by 0.05 and running the simulation again until 'stability' is reached, and continuing this process until $a = 3.5$. The red dots in Figure 4b are calculated in a similar way, but this time starting with $a = 2.05$ (just above the $a_{\max}$ bifurcation point), running to stability, then *decreasing* by 0.05 until the extent of aggregation drops to the 'no aggregation' line.

#### Supplementary Appendix I: Alternative approaches to step selection

The approach described in the Main Text, via step selection and discrete-time movement kernels, is not the only way in which researchers have attempted to scale up from animal movement processes to broad-scale space use patterns. Whilst we have deliberately focused on SSA here, we briefly mention other recent approaches.

1. **Movement as a Markov Chain Monte Carlo.** Michelot *et al.* (2018) draw an analogy between animal movements and a Monte Carlo sampling algorithm. The idea is that animals seek out a utilisation distribution by sampling the landscape as they move, rejecting locations that are less likely to correspond to the desired space use distribution.

The advantage with this approach is that the steady state UD corresponds exactly to the resource selection function (RSF) that best describes space use. However, this approach implicitly assumes that animal movement decisions emerge from a desire to construct a particular UD. If this is not the case then the resulting movement kernel may not actually reflect the animal's movement decisions. In constrast, the SSA approach attenpts to model the animal's movement decisions themselves and treats the steady state UD (if it exists) as an emergent feature of those decisions (Börger *et al.*, 2008).

2. **The Langevin equation.** Michelot *et al.* (2019) also deal with the question of describing a movement process that has an analytically-tractible UD in the form of an RSF. However, here they use a continuous time formulation, the Langevin equation, which allows for temporally irregular data.

3. **Ornstein Uhlenbeck (OU) processes.** The OU process models an animal as having some attraction towards a point, which may be fixed over all time, or may switch location depending on the environment, location, or internal state of the animal (Niu *et al.*, 2016; Wang *et al.*, 2019), as well as an inherent diffusive tendency. By incorporating landscape layers into an OU process, Eisaguirre *et al.* (2021) give a method for predicting the approximate stable home range of animals from a parametrised OU process. The method is not exact, but relies on data being sufficiently coarse-grained (Eisaguirre *et al.*, 2022).

The latter two of these use continuous time formulations, models of great research interest in recent years (Calabrese *et al.*, 2016; Niu *et al.*, 2016; Parton *et al.*, 2016; Hooten *et al.*, 2017), due conceptual and theoretical appeal, as well as their flexibility in dealing with irregular data. Formulating techniques for scaling up within a continuous time framework may ultimately be the ideal end goal. The link to PDEs is already long established in the case of OU and Langevin processes (Risken & Frank, 1996), so deserves attention in the context of animal movement. However, all the above three techniques are currently limited to situations where there is a steady-state distribution and where this depends on explanatory environmental features that are not affected by the animal locations. In other words, the effect of animal interactions, and feedbacks in general, is missing. This is an important research frontier for continuous time movement models.
